# Maternal and zygotic gene regulatory effects of endogenous RNAi pathways

**DOI:** 10.1101/453266

**Authors:** Miguel Vasconcelos Almeida, António Miguel de Jesus Domingues, René F. Ketting

## Abstract

Endogenous small RNAs (sRNAs) and Argonaute proteins are ubiquitous regulators of gene expression in germline and somatic tissues. sRNA-Argonaute complexes are often expressed in gametes and are consequently inherited by the next generation upon fertilization. In *Caenorhabditis elegans*, 26G-RNAs are primary endogenous sRNAs that trigger the expression of downstream secondary sRNAs. Two subpopulations of 26G-RNAs exist, each of which displaying strongly compartmentalized expression: one is expressed in the spermatogenic gonad and associates with the Argonautes ALG-3/4; plus another expressed in oocytes and in embryos, which associates with the Argonaute ERGO-1. The determinants and dynamics of gene silencing elicited by 26G-RNAs are largely unknown. Here, we provide diverse new insights into these endogenous sRNA pathways of *C. elegans*. Using genetics and deep sequencing, we dissect a maternal effect of the ERGO-1 branch sRNA pathway. We find that maternal primary sRNAs can trigger the production of zygotic secondary sRNAs that are able to silence targets, even in the absence of zygotic primary triggers. Thus, the interaction of maternal and zygotic sRNA populations, assures target gene silencing throughout animal development. Furthermore, we find that sRNA abundance, the pattern of origin of sRNA and 3’ UTR length are predictors of the regulatory outcome by the Argonautes ALG-3/4. Lastly, we discovered that ALG-3- and ALG-4-bound 26G-RNAs are dampening the expression of their own mRNAs, revealing a negative feedback loop. Altogether, we provide several new regulatory insights on the dynamics, target regulation and self-regulation of the endogenous RNAi pathways of *C. elegans*.

**Author Summary:** Small RNAs (sRNAs) and their partner Argonaute proteins regulate the expression of target RNAs. When sperm and egg meet upon fertilization, a diverse set of proteins and RNA, including sRNA-Argonaute complexes, is passed on to the developing progeny. Thus, these two players are important to initiate specific gene expression programs in the next generation. The nematode *Caenorhabditis elegans* expresses several classes of sRNAs. 26G-RNAs are a particular class of sRNAs that are divided into two subpopulations: one expressed in the spermatogenic gonad and another expressed in oocytes and in embryos. In this work, we describe the dynamics whereby oogenic 26G-RNAs setup gene silencing in the next generation. We also show several ways that spermatogenic 26G-RNAs and their partner Argonautes, ALG-3 and ALG-4, use to regulate their targets. Finally, we show that ALG-3 and ALG-4 are fine-tuning their own expression, a rare role of Argonaute proteins. Overall, we provide new insights into how sRNAs and Argonautes are regulating gene expression.

## Introduction

A plethora of pathways based on non-coding small RNAs (sRNAs) regulate gene expression in every domain of life. These are collectively known as RNA interference (RNAi) or RNAi-like pathways. In invertebrates, which lack adaptive immune systems and interferon response, RNAi-like pathways fulfill an immune role on the nucleic acid level, by controlling viruses and transposable elements (TEs).

MicroRNA (miRNA), Piwi-interacting RNA (piRNA) and endogenous small interfering RNA (endo-siRNA) pathways are the better described RNAi-like pathways, which differ in their biogenesis and specialized cofactors. MicroRNAs are commonly found in many, if not all, tissues and broadly regulate gene expression throughout development (1). piRNAs are typically, but not exclusively, expressed in the metazoan germline, where they assume a central function in TE control (2–5). Endo-siRNA pathways comprise varied classes of sRNAs expressed in the soma and germline that can, for example, control TEs, protein-coding genes and direct heterochromatin formation (6–8). A key commonality of RNAi-like pathways is the participation of Argonaute proteins. These proteins directly associate with sRNAs and Argonaute-sRNA complexes engage transcripts with sequence complementarity, typically resulting in target silencing. sRNA-directed gene silencing can occur both on the post-transcriptional level, by target RNA cleavage and degradation, and/or on the transcriptional level, via nuclear Argonautes that direct heterochromatin formation at target loci.

sRNAs can be viewed as genome guardians against “foreign” nucleic acids (9). In this light, the germline is an important tissue for sRNA production and function to control the transmission of “non-self” genetic elements to progeny. In multiple animals, Piwi-piRNA complexes have been shown to be maternally deposited into zygotes, where they may initiate TE silencing (10–15). Endo-siRNAs are abundantly expressed in gametes, being often required to successfully complete gametogenesis. These may also be deposited into embryos and have roles in setting up gene expression in the next generation. For example in plants, TE-derived endo-siRNAs are abundant in male and female gametes (16). Moreover, endo-siRNAs are expressed in *Drosophila* ovaries (17) and in mouse oocytes (18,19) to regulate protein-coding genes and TEs. Overall, gamete expression and maternal inheritance of Argonaute-sRNA complexes seem to be a widespread phenomenon in plants and animals, presumably important to tune gene expression during early development.

RNAi was first identified in the nematode *Caenorhabditis elegans* (20). Ever since, *C. elegans* has continuously been an important and fascinating model for studies on RNAi. *C. elegans* has an unprecedented 27 genomically encoded Argonaute genes, comprising a whole worm-specific clade of the Argonaute protein family (21). Several sRNA species have been identified in worms: miRNAs, 21U-RNAs, 22G- and 26G-RNAs (22,23). 21U-RNAs associate with PRG-1, a Piwi class Argonaute, in the germline and are therefore considered the piRNAs of *C. elegans* (24–26). 26G-RNAs can be considered primary endo-siRNAs, in that they elicit production of the overall more abundant secondary endo-siRNA pool, termed 22G-RNAs (27–29).

26G-RNAs are produced by the RNA-dependent RNA Polymerase (RdRP) RRF-3 (27–31). The ERI complex (ERIC) is an accessory complex that assists RRF-3 in producing 26G-RNAs (32–35). The conserved CHHC zinc finger protein GTSF-1 and the Tudor domain protein ERI-5 form a pre-complex with RRF-3 that is responsible for tethering the RdRP to the ERIC (32,35). Two distinct subpopulations of 26G-RNAs are synthesized in the germline and in embryos. One subpopulation is produced in the spermatogenic gonad in L4 hermaphrodites and in the male gonad, where they associate with the redundantly acting paralog Argonautes ALG-3 and ALG-4 (henceforth referred to as ALG-3/4)(27,30,31,34). These 26G-RNAs trigger the biogenesis of secondary 22G-RNAs that have been shown to either promote gene expression through the Argonaute CSR-1 or to inhibit gene expression through unidentified WAGO proteins (27,36). Hence, the effects of ALG-3/4-dependent sRNAs on their targets is complex: while some targets appear to be silenced, the expression of others seems to be positively affected. The regulatory effects resulting of the combined action of ALG-3/4 and CSR-1 seem to be more physiologically relevant at elevated temperatures (36). The conditions determining regulatory outcome, either silencing or licensing, are still unclear.

In the oogenic hermaphrodite gonad and in embryos another subpopulation of 26G-RNAs is produced. These are 3’ 2’-O-methylated by the conserved RNA methyltransferase HENN-1 (37–39) and bind to the Argonaute ERGO-1 (29). ERGO-1 targets pseudogenes, recently duplicated genes and long non-coding RNAs (lncRNAs)(29,32,40). It has recently been shown that these targets generally have a small number of introns that lack optimal splicing signals (41). ERGO-1 may thus serve as a surveillance platform to silence these inefficient transcripts, preventing detrimental accumulation of stalled spliceosomes. Effective silencing of these genes is achieved by secondary 22G-RNAs produced after ERGO-1 target recognition (28,29). In turn, these secondary 22G-RNAs may associate with cytoplasmic Argonautes that mediate post-transcriptional gene silencing (29), or to the Argonaute NRDE-3, which is shuttled into the nucleus and further silences its targets on the transcriptional level (42,43).

Depletion of spermatogenic 26G-RNAs, for example in *rrf-3, gtsf-1* and *alg-3/4* mutants, results in a range of sperm-derived fertility defects including complete sterility at higher temperatures (27,30–34). The elimination of oogenic/embryonic 26G-RNAs, for example by impairment of *rrf-3, gtsf-1* and *ergo-1*, gives rise to an Enhanced RNAi (Eri) phenotype, characterized by a response to exogenous dsRNA that is stronger than in wild-type (21,32–34). This phenotype is thought to reflect competition for common factors between exogenous and endogenous RNAi pathways (33,44). However, this Eri phenotype lacks characterization on the molecular level. Furthermore, a strong maternal rescue was reported for Eri factors (45), suggesting that maternally deposited Eri factors or their dependent sRNAs have an important role in maintaining gene silencing. The basis for this maternal rescue was not further characterized.

In this work, we address a number of gene regulatory aspects of the 26G-RNA pathways in *C. elegans*. First, we genetically dissect a maternal effect displayed by the ERGO-1 branch of the 26G-RNA pathway. Our findings suggest that both maternal and zygotic sRNAs drive gene silencing throughout embryogenesis and larval development until adulthood. Furthermore, we explore ALG-3/4 target regulation and find that sRNA abundance, origin of the sRNAs and 3’ UTR length are predictors of the regulatory outcome. Lastly, we find that the 26G-RNA-binding Argonautes ALG-3 and ALG-4 negatively regulate their own expression, which, to our knowledge, represents the first description of such regulatory feedback mechanism amongst *C. elegans* Argonautes.

## Results

### Maternal and zygotic endogenous small RNAs drive RNAi in the soma

*rrf-3* and *gtsf-1* mutants lack the two subpopulations of 26G-RNAs, and display the phenotypes associated with depletion of both subpopulations: the enhanced RNAi (Eri) phenotype, shared with *ergo-1* mutants (21,32–34), and sperm-derived fertility defects, shared with *alg-3/4* double mutants (27,30–34,36). **S1A Fig** offers a simplified scheme of these pathways. For clarity, the two subpopulations of 26G-RNAs and downstream 22G-RNAs, dependent on ERGO-1 or ALG-3/4 will be referred to as ERGO-1 branch sRNAs and ALG-3/4 branch sRNAs, respectively.

We have previously shown that germline-specific GTSF-1 transgenes could rescue the enhanced RNAi (Eri) phenotype of *gtsf-1* mutants (32). This was an intriguing result, since the Eri phenotype arises after targeting somatically expressed genes with RNAi, indicating that germline-expressed GTSF-1 is able to affect RNAi in the soma, possibly through maternal deposition of GTSF-1 or GTSF-1-dependent sRNAs. We reasoned that if maternal GTSF-1 activity can prime gene silencing in embryos then the transmission of the Eri phenotype should show a maternal rescue. To address this experimentally, we linked *gtsf-1(xf43)* to *dpy-4(e1166)*, and crossed the resulting double mutants with wild-type males (**Fig 1A**). We then allowed for two generations of heterozygosity and assayed for RNAi sensitivity in homozygous *gtsf-1* mutant F1 and F2 generations, scoring for larval arrest triggered by *lir-1* RNAi. Indeed, the Eri phenotype showed a strong maternal effect, arising only in the F2 generation of *gtsf-1* mutants (**Fig 1A**). This is consistent with a maternal effect reported for other Eri factors (45). We have previously shown that GTSF-1 is required to silence a GFP transgene reporting on ERGO-1 branch 22G-RNA activity, referred to as 22G sensor (32). Therefore, we also looked at the dynamics of derepression of this transgene upon introduction of *gtsf-1* mutation. We noticed that strong GFP expression appeared only in the second generation of homozygosity of the *gtsf-1* allele (**Fig 1B-C**). An identical maternal effect on the expression status of this transgene is observed after crossing in *rrf-3, ergo-1* and other *gtsf-1* mutant alleles (**S1B Fig**). Combined with our previously described rescue of the Eri phenotype using a germline promoter, these results strongly suggest that maternally provided ERGO-1 branch pathway components are sufficient to establish normal RNAi sensitivity in the soma of *C. elegans*.

**Fig 1.**
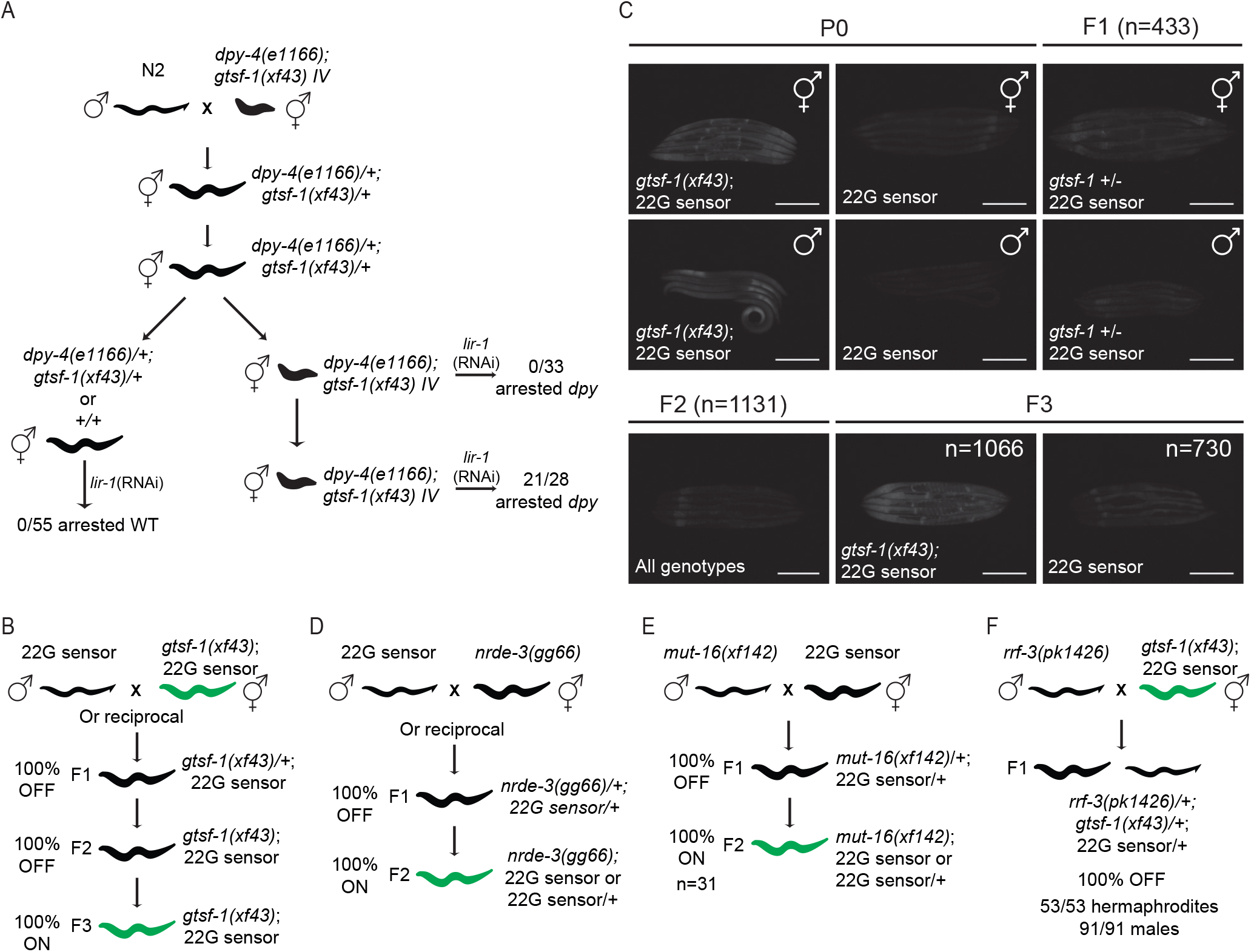
**Maternal and zygotic sRNAs drive RNAi in the soma**. (A) Experimental setup to address maternal transmission of the Eri phenotype in *gtsf-1* mutants. Eri phenotype was assessed by transferring worms to plates containing *lir-1* RNAi food and scoring for larval arrest. *dpy-4(e1166)* is weakly semi-dominant. Since the phenotype is mild, for simplicity, we will refer to *dpy-4(e1166)* heterozygotes as “wild-type”. (B, D-F) Schematics of genetic crosses using the 22G sensor background. Green worms illustrate ubiquitous derepression of 22G sensor. Unless otherwise noted, for all crosses the number of scored F1s, F2s and F3s was each >50. (C) Related to the cross shown in (B). Wide-field fluorescence microscopy images showing 22G sensor GFP signal. Five representative gravid adult hermaphrodites or adult males from each generation are shown. Of note, some autofluorescence of the gut is observed in gravid adult animals, and is especially noticeable in worms with the sensor off. Scale bars represent 0,25 mm.

Although the silencing of the 22G sensor used in our experiments is dependent on ERGO-1, ERGO-1 is not the Argonaute protein binding to the effector 22G-RNA (29,39). This has been shown to be driven by the somatically expressed, nuclear Argonaute protein NRDE-3 (40,42), and maybe additional cytoplasmic WAGOs (29)(**S1A Fig**). In absence of ERGO-1 and other 26G-RNA pathway factors, NRDE-3 is no longer nuclear, and in *nrde-3* mutants the 22G sensor is activated, indicating that NRDE-3 requires sRNA input from ERGO-1 branch sRNAs (32,39,40,42). Strikingly, loss of NRDE-3 derepressed the 22G sensor transgene in the first homozygous generation (**Fig 1D**), showing that in contrast to 26G-RNAs, the downstream 22G-RNA pathway is not maternally provided. MUT-16 is a factor required for the nucleation of mutator foci and 22G-RNA biogenesis (46). Confirming the requirement for zygotically produced 22G-RNAs, absence of MUT-16 derepresses the 22G sensor in the first homozygous mutant generation (**Fig 1E**). These results suggest a scenario in which 1) NRDE-3 is loaded with zygotically produced 22G-RNAs that are primed by maternally provided 26G-RNAs and 2) NRDE-3 activity is maintained in somatic tissues until the adult stage, in absence of a zygotic 26G-RNA pathway.

The results presented above show that maternal 26G-RNAs are sufficient for 22G sensor silencing. We also tested whether maternal 26G-RNAs are necessary for 22G sensor silencing by crossing *rrf-3* mutant males with *gtsf-1; 22G sensor* hermaphrodites (**Fig 1F**). Both of these strains lack 26G-RNAs and their downstream 22G-RNAs, therefore, their progeny will not receive a maternal and/or paternal complement of these sRNAs. The 22G sensor was silenced in all cross progeny, showing that in the absence of maternal 26G-RNAs, zygotic 26G-RNAs can induce production of silencing-competent 22G-RNAs. Thus, maternal 26G-RNAs appear to be sufficient but not necessary for target silencing.

### 26G-RNA-derived maternal effects are restricted to the ERGO-1 branch

The maternal effects described above for the Eri phenotype and for 22G sensor silencing, are related to the ERGO-1 branch of the pathway. Next, we wanted to determine if the ALG-3/4 branch also displays such a parental effect. To test this, we assessed the influence of maternal GTSF-1 activity on the temperature-sensitive sterility phenotype. Using the same setup as we used for the Eri experiment (in **Fig 1A**), we observe that the temperature-sensitive sperm defect of *gtsf-1* mutants was not rescued maternally (**S1C Fig**), indicating that such maternal effects are likely restricted to the ERGO-1 branch.

### Maternal GTSF-1 supports zygotic ERGO-1 branch 22G-RNA production

The 22G sensor reports on the silencing activity of a single 22G-RNA that maps to the so-called X-cluster, a known set of targets of ERGO-1 (29,39). Therefore, the experiments above using this 22G sensor have a limited resolution and our observations may not reflect the silencing status of most ERGO-1 targets. To characterize this maternal effect in more detail and in a broader set of ERGO-1 targets, we decided to analyze sRNA populations in young adult animals. Concretely, we outcrossed *dpy-4*; *gtsf-1* and sequenced sRNAs from wild-type and two consecutive generations of Dpy young adult animals (**Fig 2A**). First generation *gtsf-1* homozygous mutants will henceforth be addressed as “mutant F1”, and second generation *gtsf-1* homozygous mutants as “mutant F2” (**Fig 2A**). We sequenced young adult animals because they lack embryos, therefore avoiding confounding effects with zygotic sRNAs of the next generation. sRNAs were cloned and sequenced from four biological replicates. The cloning of sRNAs was done either directly (henceforth referred to as untreated samples) or after treatment with the pyrophosphatase RppH (47) before library preparation. The latter enriches for 22G-RNA species that bear a 5’ triphosphate group. Sequenced sRNAs were normalized to all mapped reads excluding structural reads (sequencing statistics can be found in the **S1 Table**). In our analysis we strictly looked at 26G- and 22G-RNAs that map in antisense orientation to protein-coding and non-coding genes (see **Methods**).

**Fig 2.**
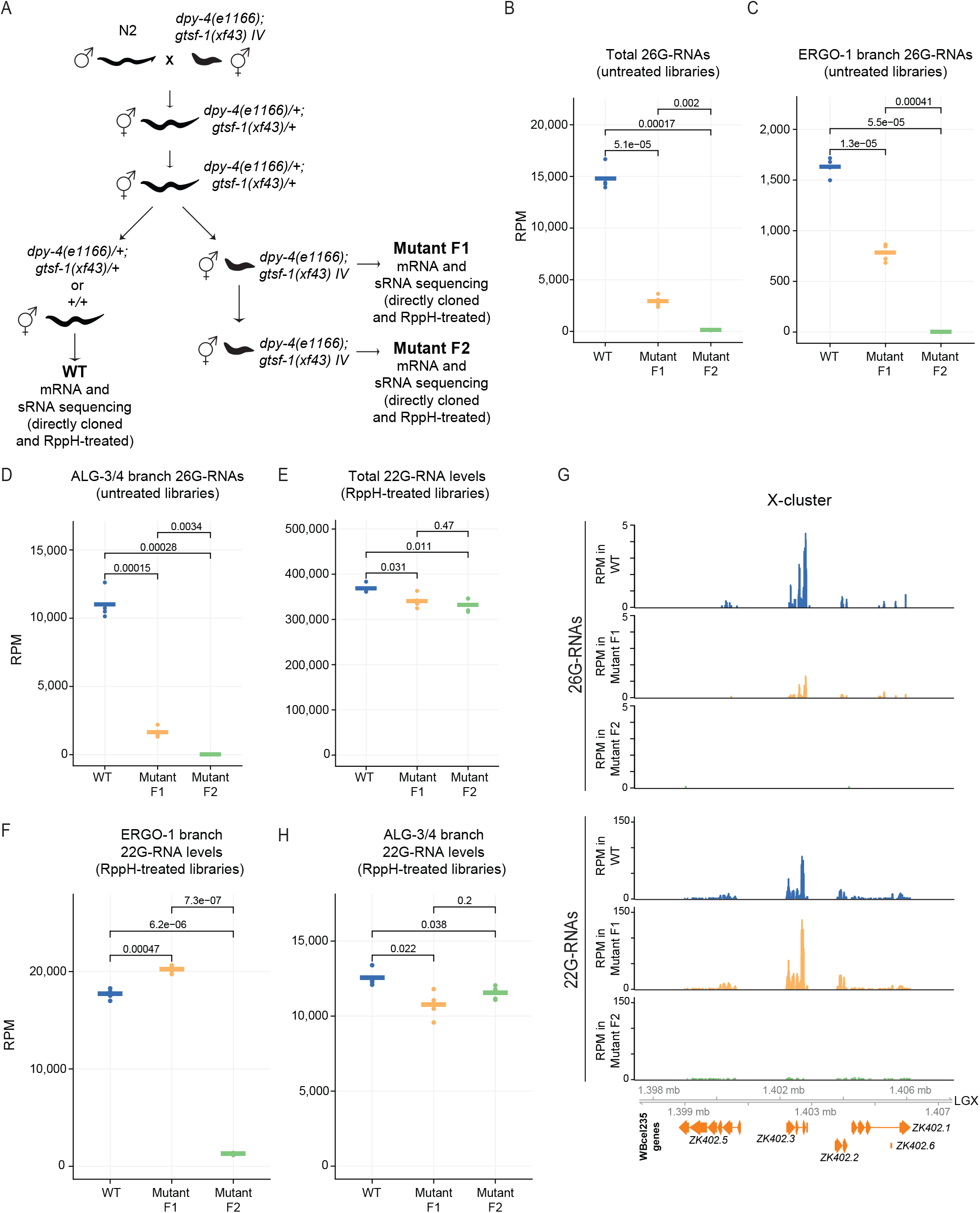
**sRNA dynamics in Eri maternal inheritance**. (A) Schematics of the cross setup used to isolate worms of different generations and *gtsf-1* genotypes. *gtsf-1;dpy-4* mutants were outcrossed with N2 males, allowed to self for two generations and then WT, Dpy mutant F1 and Dpy mutant F2 young adult animals were isolated, RNA was extracted, sRNAs and mRNAs were cloned and sequenced. sRNA libraries were either prepared directly or after treatment with RppH. WT, wild-type. (B-F, H) Normalized levels of sRNAs, in RPM (Reads Per Million), per generation/phenotype. Four biological replicates are shown. (B) Total levels of 26G-RNAs in the untreated libraries. (C) Total levels of 26G-RNAs mapping to ERGO-1 targets in the untreated libraries. (D) Total levels of 26G-RNAs mapping to ALG-3/4 targets in the untreated libraries. (E) Total levels of 22G-RNAs in the RppH-treated libraries. (F) Total levels of 22G-RNAs mapping to ERGO-1 targets in the RppH-treated libraries. (G) Genome browser tracks of the X-cluster, a known set of ERGO-1 targets, showing mapped 26G- and 22G-RNAs. 26G- and 22G-RNA tracks were obtained from untreated and RppH-treated libraries, respectively. (H) Total levels of 22G-RNAs mapping to ALG-3/4 targets in RppH-treated libraries. P-values were calculated with a two-sided unpaired t-test.

Total 26G-RNA levels are depleted in young adults lacking GTSF-1 (**Fig 2B**). Mutant F1s have significantly less 26G-RNAs than wild-type worms, while mutant F2s have 26G-RNA levels very close to zero (**Fig 2B**). For a finer analysis we looked specifically at 26G-RNAs derived from ERGO-1 and ALG-3/4 targets (as defined in reference 32, see **Methods**). 26G-RNAs mapping to these two sets of targets recapitulate the pattern observed for global 26G-RNAs (**Fig 2C-D**). The difference between the F1 and F2 mutants might reflect a maternal 26G-RNA pool that is still detectable in the young adult F1, but no longer in the F2. However, we point out that amongst the selected F1 Dpy animals, approximately 5.2% will in fact be *gtsf-1* heterozygous, due to meiotic recombination between *gtsf-1* and *dpy-4* (estimated genetic distance between these two genes is 2.6 map units). Hence, another explanation for the mutant F1 pool of 26G-RNAs may be a contamination of the *gtsf-1* homozygous pool with heterozygous animals. The mutant F2 was isolated from genotyped F1 animals, excluding this confounding effect. We conclude that in young adult mutant F1 animals, maternally provided 26G-RNAs (or 26G-RNAs produced zygotically by maternal proteins) are no longer detectable at significant levels.

Total levels of 22G-RNAs are slightly reduced in mutant F1 and F2 animals (**Fig 2E**). However, total 22G-RNA levels encompass several distinct subpopulations of 22G-RNAs, including those that do not depend on 26G-RNAs. To have a closer look on 22G-RNAs that are dependent on 26G-RNAs, we focused on 22G-RNAs that map to ERGO-1 and ALG-3/4 targets. Strikingly, compared to wild-type, the 22G-RNA population from ERGO-1 targets is moderately higher in mutant F1 animals, and are subsequently depleted in the mutant F2 generation (**Figs 2F and S2A**). These effects are not only clear in overall analysis, but also on a well-established set of ERGO-1 branch targets, such as the X-cluster (**Fig 2G**). Consistent with a role of NRDE-3 downstream of ERGO-1, 22G-RNAs mapping to annotated NRDE-3 targets (43) show the same pattern of depletion as ERGO-1-dependent 22G-RNAs (**S2A Fig**). These results are consistent with the idea that the Eri phenotype and 22G sensor derepression are caused by the absence of NRDE-3-bound, secondary 22G-RNAs downstream of 26G-RNAs.

ALG-3/4-dependent downstream 22G-RNAs behave differently in this experiment (**Fig 2H**). Upon disruption of *gtsf-1*, ALG-3/4-dependent 22G-RNAs are only slightly affected in both the mutant F1 and F2 (**Figs 2H and S2A**), despite the fact that their upstream 26G-RNAs are absent. This is illustrated in **S2B Fig** with genome browser tracks of *ssp-16*, a known ALG-3/4 target. We conclude that 26G-RNA-independent mechanisms are in place to drive 22G-RNA production from these genes.

Finally, 21U-RNAs and 22G-RNAs mapping to other known RNAi targets are not affected in this inheritance setup, supporting the notion that *gtsf-1* is not affecting these sRNA species (**S2A, C Fig**). One exception are the 22G-RNAs from CSR-1 targets, which seem to be slightly depleted in both the mutant F1 and F2 generations (**S2A Fig**). It is not possible to dissect whether this is a direct effect or not, but we note that mRNA levels of CSR-1 targets are slightly downregulated in the analyzed mutants (**S2D Fig**). Given that CSR-1 22G-RNAs tend to correlate positively with gene expression (48), it is conceivable that the reduction of CSR-1 target 22G-RNAs is the result of decreased target gene expression.

### ERGO-1 pathway mRNA targets show stronger upregulation in the second *gtsf-1* homozygous mutant generation

The very same samples used for generating sRNA sequencing data were also used for mRNA sequencing (**Fig 2A**). First, we checked *gtsf-1* expression. As expected, *gtsf-1* is strongly depleted in the mutant samples (**S3A Fig**). In the mutant F1 we still observe a low level of *gtsf-1* derived transcripts (about 9.5% of wild-type) that is absent from the mutant F2. These transcripts cover the region deleted in the *gtsf-1(xf43)* mutant allele, indicating they cannot represent zygotically transcribed *gtsf-1* mutant mRNA. Rather, they likely come from the above described contamination of the homozygous F1 population with heterozygous animals.

We hypothesized that ERGO-1 branch 22G-RNAs observed in the mutant F1 generation might be competent to maintain target silencing. If this is true, we should observe strong upregulation of ERGO-1 target mRNAs only the mutant F2 generation. Indeed, the X-cluster is upregulated only in the second mutant generation (**Fig 3A**). When ERGO-1 targets are analyzed in bulk, we observe the same trend, with stronger upregulation only in the mutant F2, consistent with the maternal effect (**Fig 3B**). ALG-3/4 targets, as for instance *ssp-16*, were found to be upregulated already in the F1 generation (**Figs 3B** and **S3B**), supporting the notion that the maternal rescue of the 26G-RNA pathways is restricted to the ERGO-1 branch.

**Fig 3.**
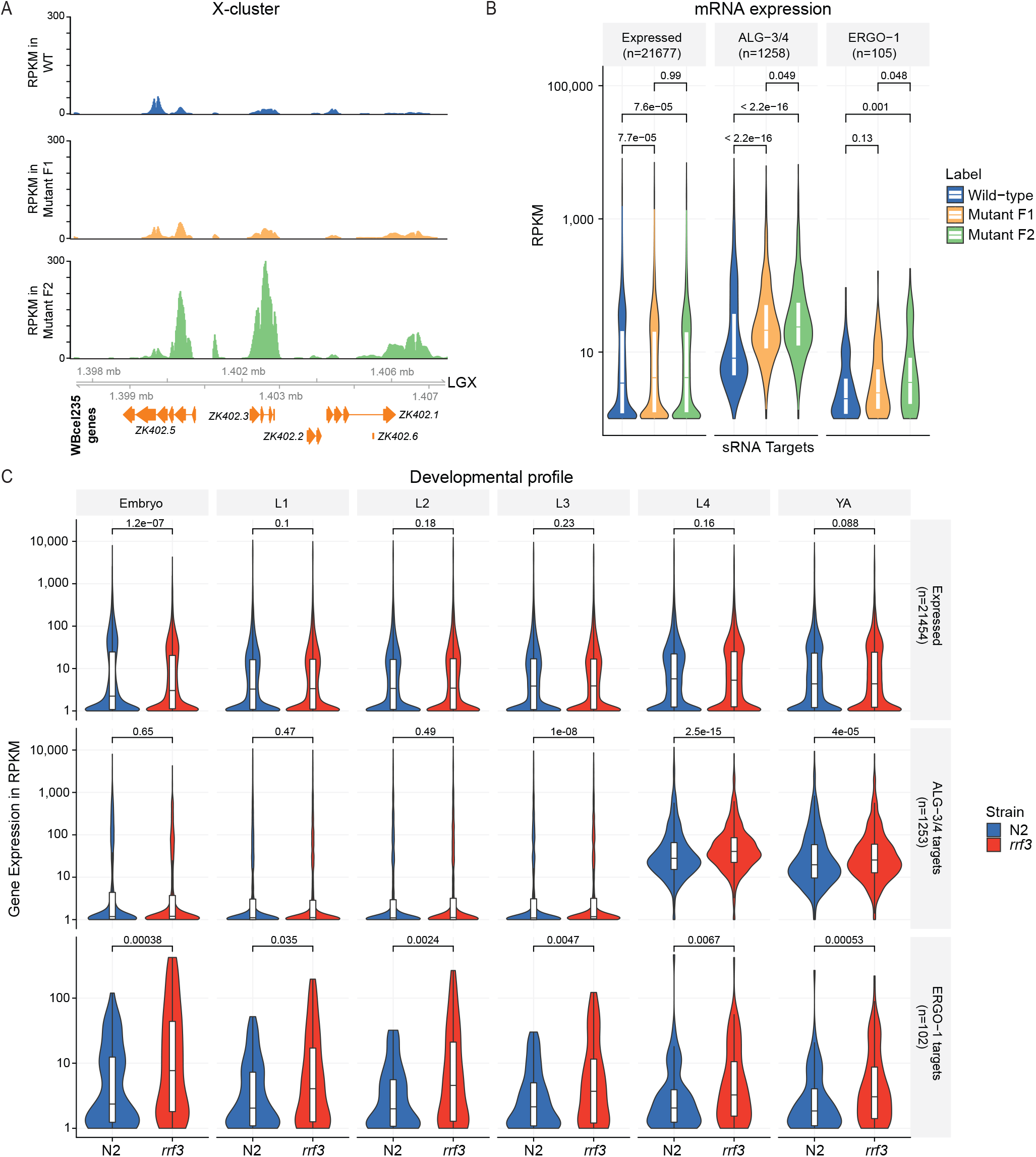
**mRNA dynamics in Eri maternal inheritance**. (A) Genome browser tracks showing mRNA, in RPKM (Reads Per Kilobase Million), of X-cluster genes. (B) Distribution of normalized mRNA expression, in RPKM, of all expressed genes, ALG-3/4 targets and ERGO-1 targets in different generations/phenotype. (C) Distribution of normalized mRNA expression, in RPKM, of all expressed genes (upper panel), ALG-3/4 targets (middle panel) and ERGO-1 targets (lower panel) throughout development. Expression is shown for wild-type N2 (in blue) and *rrf-3(pk1426)* (in red) animals. YA, young adult. L1-L4, first to fourth larval stages of *C. elegans* development. Violin plots in (B-C) show the distribution density of the underlying data. The top and bottom of the embedded box represent the 75th and the 25th percentile of the distribution, respectively. The line in the box represents the median. P-values were calculated with a two-sided unpaired Mann-Whitney/Wilcoxon rank-sum test.

### Eri targets show stronger expression in embryos

ERGO-1 targets comprise a very diverse set of targets consisting of pseudogenes, fast evolving small genes, paralog genes and lncRNAs (29,40,41). Considering the maternal effect described above for ERGO-1-dependent sRNA and correspondent target, we postulated that this maternal effect may exist to counteract embryonic expression of ERGO-1 targets. To address this we sequenced mRNA of synchronized populations of all developmental stages (L1, L2, L3, L4, young adult and embryos) of both wild-type (N2) and *rrf-3(pk1426)* mutants. Global gene expression in *rrf-3* mutants is significantly different only in embryos (**Fig 3C**, upper panel). These changes may be explained by higher expression of ERGO-1 targets during embryogenesis. Indeed, in wild-type worms, ERGO-1 targets are most abundant in embryos (**Fig 3C**, lower panel, in blue). Moreover, the effect of *rrf-3* mutation on ERGO-1 target expression is stronger in embryos (**Fig 3C**, lower panel). These results indicate that the maternal effect reported above can reflect deposition of factors which are required to initiate silencing of targets early in development.

### GTSF-1 is required for sRNA biogenesis and target silencing in the male germline

The young adult sequencing datasets we obtained in this study (**Fig 2A**), as well as previous datasets of gravid adults (32), are not well suited to address ALG-3/4 biology, considering that in these developmental stages ALG-3/4 are not expressed, at least not abundantly. Therefore, in order to further our understanding of the dependency of ALG-3/4 branch sRNAs on GTSF-1, we generated additional sRNA and mRNA datasets from wild-type and *gtsf-1* male animals grown at 20°C.

As expected, global 26G-RNA levels are severely affected in *gtsf-1* mutant males, reflecting downregulation of 26G-RNAs from both branches of the pathway (**Fig 4A-C**). Consistent with the absence of ERGO-1 in adult males, ERGO-1 branch 26G-RNAs are detected in extremely low numbers in wild-type animals (**Fig 4B**). Global levels of 21U-RNAs seem to be moderately augmented (**Fig 4D**), possibly resulting from the lack of 26G-RNAs in the libraries. Global levels of 22G-RNAs are not affected (**Fig 4E**), but consistent with a global depletion of 26G-RNAs, 22G-RNAs specifically mapping to ALG-3/4 and ERGO-1 targets are reduced in *gtsf-1* mutant males (**Fig 4F**). Next, we probed the effects of *gtsf-1* mutation on male gene expression using mRNA sequencing. ALG-3/4 and ERGO-1 targets are both upregulated *gtsf-1* mutant males (**Fig 4G**). These changes are illustrated for the X-cluster and *ssp-16* in the genome browser tracks of **S4 Fig**.

**Fig 4.**
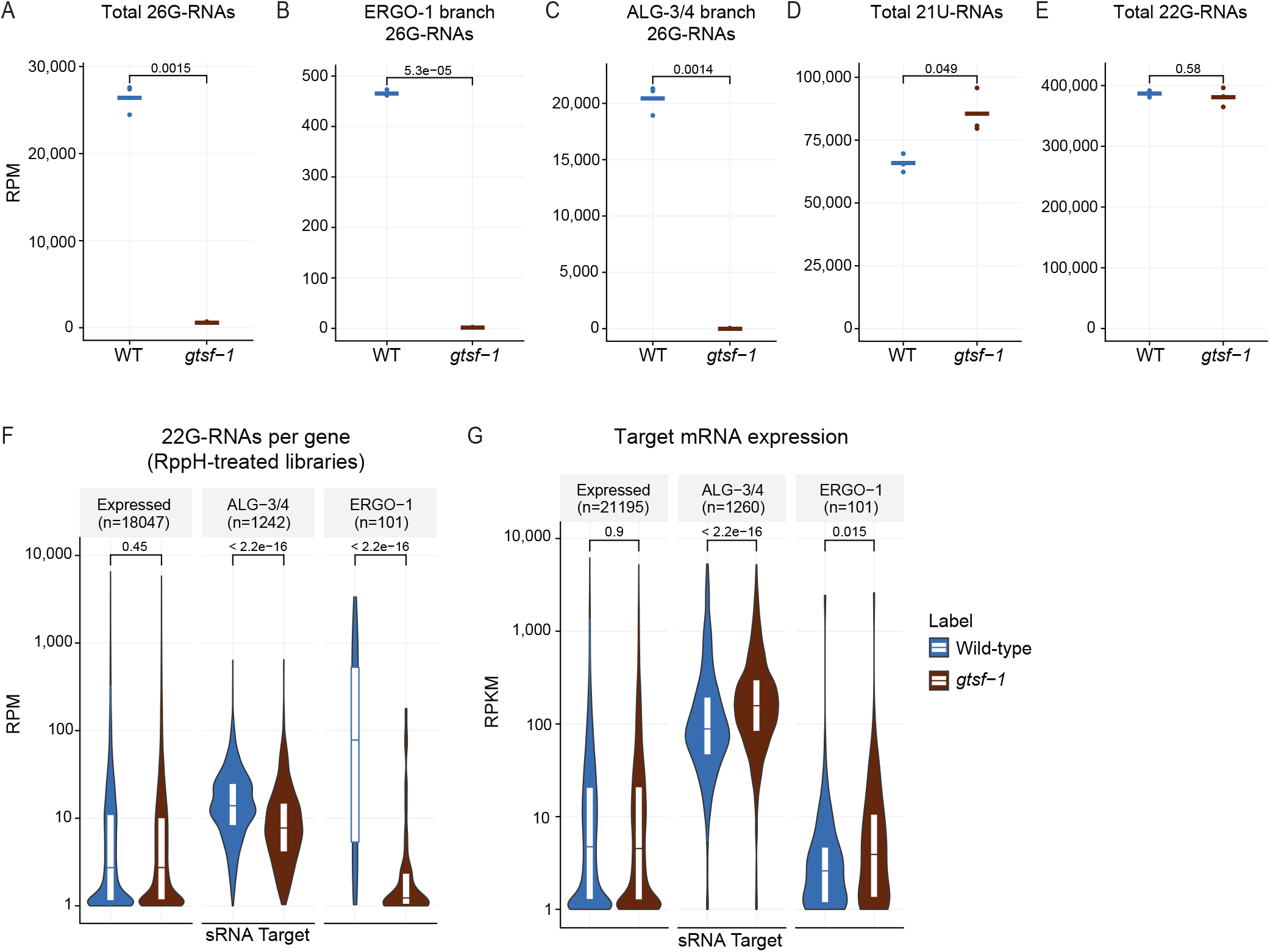
**GTSF-1 is required for sRNA biogenesis and target silencing in adult males**. (A-E) Normalized levels of sRNAs in RppH treated libraries, in RPM. Three biological replicates are shown. WT, wild-type. (A) Total levels of 26G-RNAs. (B) 26G-RNAs mapping to ERGO-1 targets. (C) 26G-RNAs mapping to ALG-3/4 targets. (D) total levels of 21U-RNAs. (E) total levels of 22G-RNAs. (F) RPM Levels of sRNAs mapping, per gene, to known targets of ALG-3/4 and ERGO-1. (G) Normalized mRNA expression of ALG-3/4 and ERGO-1 targets, in RPKM. Violin plots in (F-G) show the distribution density of the underlying data. The top and bottom of the embedded box represent the 75th and the 25th percentile of the distribution, respectively. The line in the box represents the median. P-values were calculated either with a two-sided unpaired t-test (A-E), or with a two-sided Mann-Whitney/Wilcoxon rank-sum test (F-G).

As a final note on the developmental aspects of ALG-3/4 branch, consistent with enrichment in the spermatogenic gonad (27,30–32,34,36), ALG-3/4 targets are more highly expressed and more responsive to *rrf-3* mutation in the L4 and young adult stages of hermaphrodite animals (**Fig 3C**, middle panel). Given that the overall ALG-3/4 target mRNA levels go up upon depletion of *gtsf-1* or *rrf-3* (**Figs 3C** and **4G**), bulk 26G-RNA activity during spermatogenesis seems to be repressive at 20°C.

We conclude that the activity of GTSF-1 is required in the male germline for silencing of 26G-RNA targets by participating in 26G- and 22G-RNA biogenesis.

### 22G-RNA abundance is a predictor of the regulatory outcome of ALG-3/4 targets

ALG-3/4 were shown to have distinct effects on gene expression, either silencing or licensing (27,36). However, how these different effects arise is currently unknown. Even though our analysis in males did not reveal a licensing effect of 26G RNAs, the bulk analysis of targets in **Figs 3B** and **4G** may occlude the behavior of distinct target subpopulations. Of note, our sequencing datasets were obtained from animals grown at 20°C and are therefore blind to the strong positive regulatory effect of ALG-3/4 in gene expression at higher temperatures (36).

We reasoned that sRNA abundance may be correlated with different regulatory outcomes. Therefore, we defined ALG-3/4 targets that are upregulated, downregulated and unaltered upon *gtsf-1* mutation and plotted their 26G-RNA abundance. This reveals a tendency for genes that are upregulated upon loss of GTSF-1 to be more heavily targeted by 26G-RNAs in the adult male germline (**Fig 5A**, left panel). The same trend is observed for 22G-RNAs: upregulated genes are more heavily covered by 22G-RNAs (**Fig 5A**, right panel, and **5B**). In contrast, ALG-3/4 targets that are downregulated in *gtsf-1* mutant males display a relatively low-level targeting by 22G-RNAs (**Fig 5A-B**).

**Fig 5.**
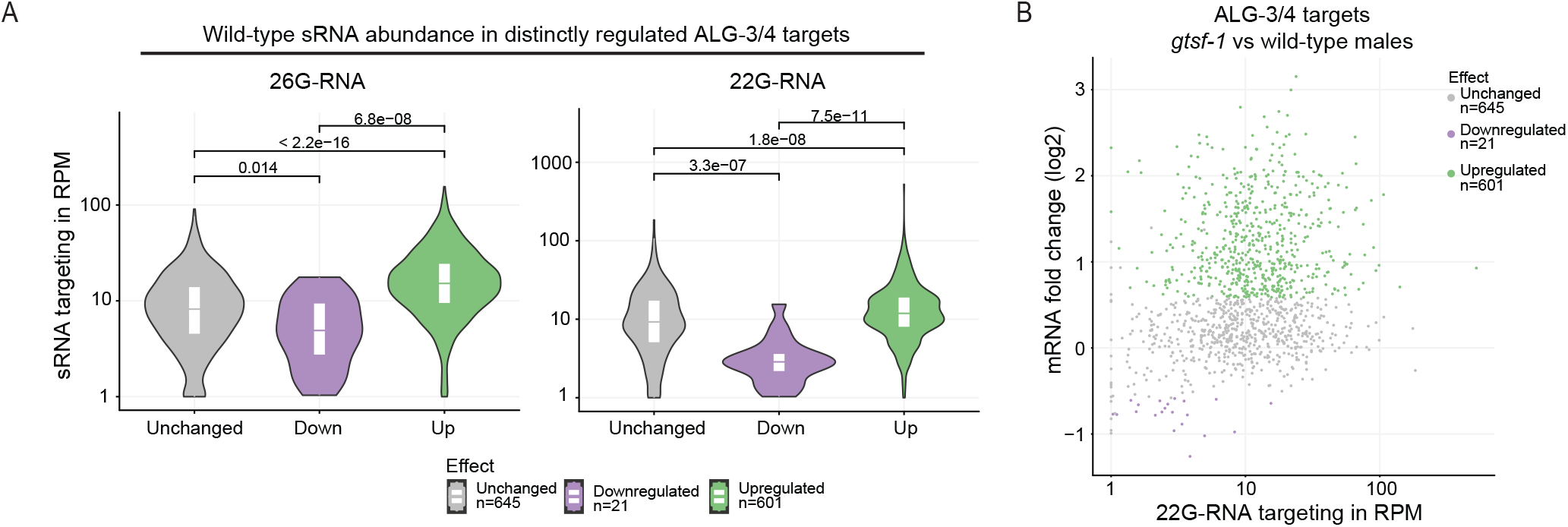
**sRNA abundance is a predictor of regulatory outcome by ALG-3/4**. (A) Distribution of sRNA levels (26G-RNA on the left panel, 22G-RNA on the right panel) mapping to ALG-3/4 targets that are unchanged, down- or upregulated upon *gtsf-1* mutation. (B) MA-plot displaying the 22G-RNA levels in respect to regulatory outcome. (B) is another representation of the data shown in the right panel of (A). Violin plots in (A) show the distribution density of the underlying data. The top and bottom of the embedded box represent the 75th and the 25th percentile of the distribution, respectively. The line in the box represents the median. P-values were calculated with a two-sided unpaired Mann-Whitney/Wilcoxon rank-sum test.

We conclude that stronger 26G-RNA targeting promotes stronger 22G-RNA biogenesis and repression of targets, whereas low-level targeting by 26G- and 22G-RNAs does not. Transcripts that are downregulated in absence of GTSF-1 might be licensed for gene expression, but may also respond in a secondary manner to a disturbed 26G-RNA pathway.

### ALG-3/4- and ERGO-1-branch 26G-RNA subpopulations display different patterns of origin that influence target expression

It was previously noticed that ALG-3/4-dependent 26G-RNAs mostly map to both the 5’ and 3’ ends of their targets, and that this may correlate with gene expression changes (27). We followed up on this observation by performing metagene analysis of 26G-RNA binding using our broader set of targets. Indeed, ALG-3/4 branch 26G-RNAs display a distinctive pattern with two sharp peaks near the transcription start site (TSS) and transcription end site (TES) (**Figs 6A** and **S5A**, left panels). In contrast, ERGO-1 branch 26G-RNAs map throughout the transcript, with a slight enrichment in the 3’ half (**Figs 6B**, left panel). Contrary to 26G-RNAs, 22G-RNAs from both branches map throughout the transcript (**Figs 6A-B** and **S5A**, right panels). These patterns are consistent with recruitment of RdRPs and production of antisense sRNAs along the full length of the transcript. These findings suggest substantially different regulation modes by ERGO-1- and ALG-3/4-branch 26G-RNAs.

**Fig 6.**
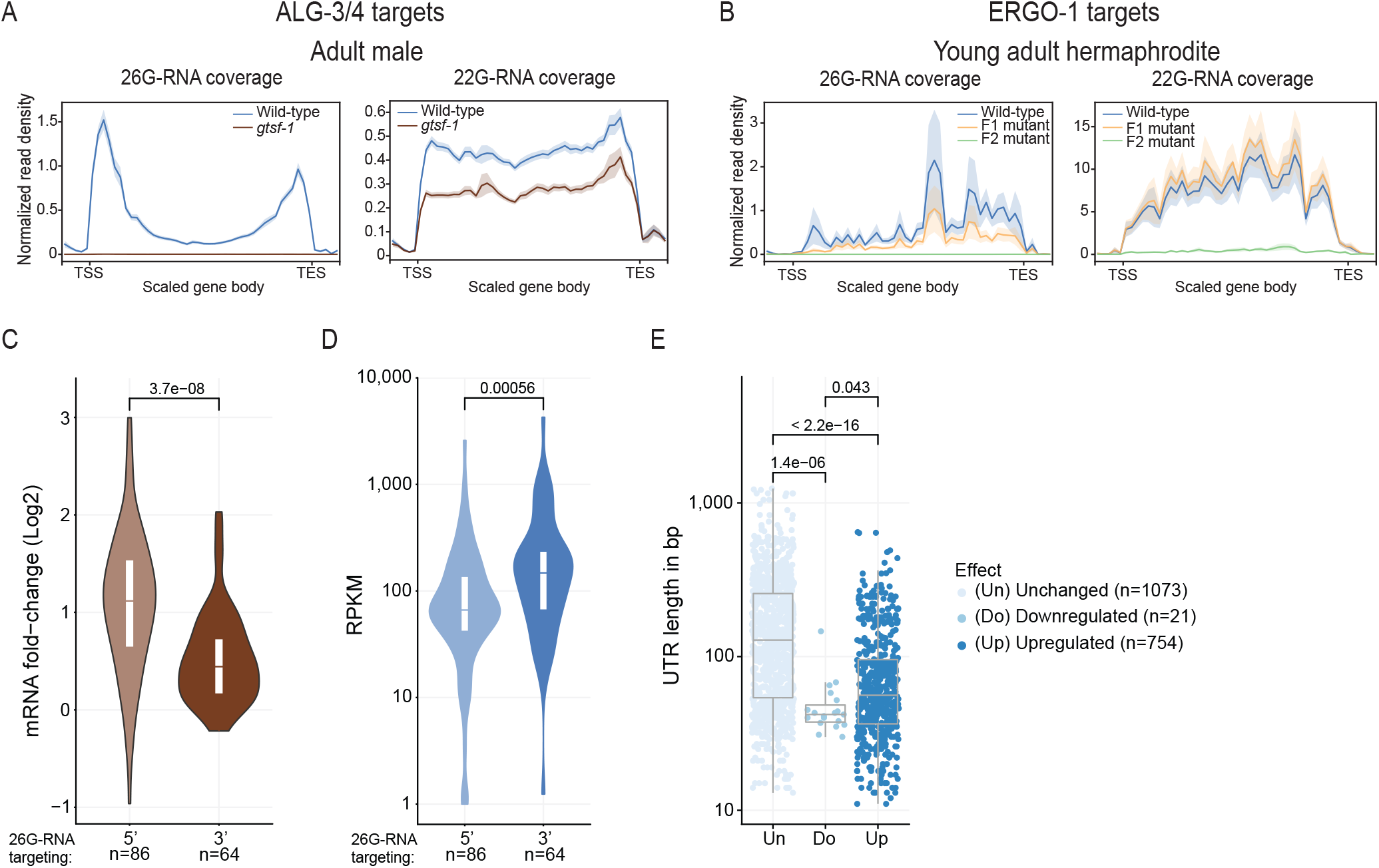
**Predictors of regulatory outcome by ALG-3/4 in males, and ERGO-1 branch sRNA metagene analysis**. (A-B) Metagene analysis of 26G- (left panel) and 22G-RNAs (right panel) mapping to ALG-3/4 targets (n=1258) in male datasets (A), and to ERGO-1 targets (n=104) in young adult datasets (B), from our maternal effect setup (as in **Fig 2A**). Target gene body length was scaled between transcription start site (TSS) and transcription end site (TES). Moreover, the regions comprising 250 nucleotides immediately upstream of the TSS and downstream of the TES are also included. (C) Regulation of ALG-3/4 target genes predominantly targeted on the 5’ or on the 3’ by 26G-RNAs. (D) Wild-type expression levels, in RPKM, of ALG-3/4 target genes predominantly targeted on the 5’ on or the 3’ by 26G-RNAs. (E) 3’ UTR lengths of all the transcript isoforms annotated for ALG-3/4 target genes, according to effect on gene expression. For (C-E) we used male sequencing datasets. Violin plots in (C-D) and the boxplot in (E) show the distribution of the data. The top and bottom of the embedded boxes represent the 75th and the 25th percentile of the distribution, respectively. The line in the box represents the median. P-values were calculated with a two-sided unpaired Mann-Whitney/Wilcoxon rank-sum test.

Conine and colleagues reported a correlation between 26G-RNA 5‘targeting and negative regulation (27). We wanted to address whether our datasets show concrete correlations between the patterns of origin of ALG-3/4-dependent 26G-RNAs and distinct regulatory outcomes. To address this, we ranked genes by 5’ and 3’ abundance of 26G-RNAs, selected genes predominantly targeted at the 5’ or on the 3′ ends and plotted their fold change upon *gtsf-1* mutation. Dominant 5’ targeting by 26G-RNAs seems to be correlated with gene silencing (fold change >0 in the mutant, **Fig 6C**), whereas dominant 3’ targeting is accompanied with only weak upregulation in *gtsf-1* mutant males (**Fig 6C**). In further support for a non-gene silencing, and potentially licensing role for ALG-3/4 targeting the 3’ end, genes with predominant 3’ 26G-RNAs display an overall higher expression that genes predominantly targeted on the 5’ region (**Fig 6D**). The same signatures are found in young adults, with an even stronger signature of the 3’ in promoting gene expression (**S5B-C Fig**).

Finally, we interrogated if the length of 5’ and 3’ UTRs may be a predictor of regulatory outcome by ALG-3/4. 5’ UTR length was not significantly different between unchanged, downregulated and upregulated genes (unpublished observations). In contrast, 3’ UTR length is significantly smaller in targets that respond to loss of GTSF-1 in males (**Fig 6E**). Interestingly, we find the same and possibly even stronger relation between 3’UTR length and responsiveness to GTSF-1 status in young adult animals (**S5D Fig**).

Altogether, our results suggest that in males, 3’ vs 5’ targeting and 3’ UTR length are predictors of whether ALG-3/4 targets are silenced or not.

### ALG-3 and ALG-4 act in a negative feedback loop

While navigating the lists of GTSF-1 targets defined by differential gene expression analysis, we noticed that *alg-3* and *alg-4* are targets of 26G-RNAs (this study and in reference 32). These 26G-RNAs are sensitive to oxidation (not enriched in oxidized libraries, see reference 32) and map predominantly to the extremities of the transcript (**Figs 7**, upper panels), indicating that these 26G-RNAs share features with ALG-3/4 branch 26G-RNAs. In addition to these 26G-RNAs, significant amounts of 22G-RNAs are found on *alg-3/4* (**Fig 7**, middle panels). These sRNAs seem to silence gene expression, since mRNA-seq shows that *alg-3* and *alg-4* transcripts are 2-3 fold upregulated in *gtsf-1* mutants (**Fig 7**, lower panels).

**Fig 7.**
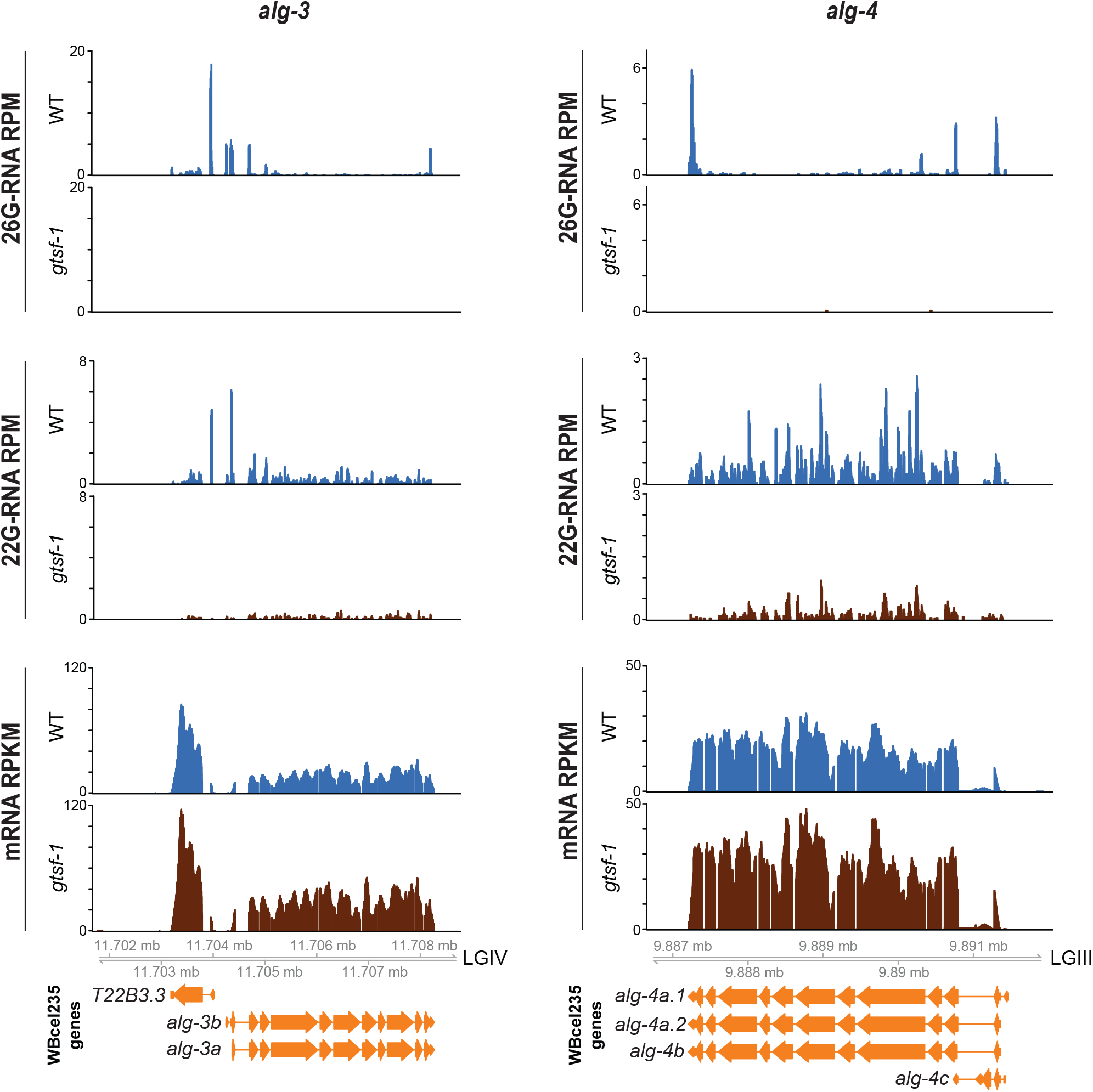
**ALG-3 and ALG-4 are engaged in a negative feedback loop in males**. Genome browser tracks showing 26G-RNAs (upper panels) and 22G-RNAs (middle panels) mapping to *alg-3* (left panels) and *alg-4* (right panels), in RPM. Lower panels show the RPKM mRNA levels of *alg-3* (on the left) and *alg-4* (on the right). Sequencing datasets of adult males were used. WT, wild-type.

These results strongly suggest that *alg-3/4* are regulating their own expression in a negative feedback loop. Of note, the upregulation of *alg-3* and *alg-4* is in agreement with the results presented above, because these genes are more heavily targeted by 26G-RNAs on their 5’ (although *alg-4* also has a sharp 3’ 26G-RNA peak, upper panels). Furthermore, these same signatures of negative feedback loop are observed in young adults (**S6 Fig**).

## Discussion

### Genetic dissection of a maternal rescue

Animal male and female gametes are rich in RNA. Upon fertilization, several RNA species are thus provided to the zygote. Multiple lines of evidence from several distinct organisms indicate that sRNAs are included in the parental repertoire of inherited RNA. For example, piRNAs have been reported to be maternally deposited in embryos in *D. melanogaster* and *C. elegans* (10,12,13,15,49). In *C. elegans* other endogenous sRNA populations have also been shown to be contributed by the gametes: 1) 26G-RNAs have been shown to be weakly provided by the male, while 22G-RNAs are more abundantly provided (50); 2) 26G-RNAs and the Argonaute ERGO-1 are co-expressed during oogenesis and in embryos (29,31,51); and 3) 22G-RNAs are deposited in embryos via the mother and participate in transgenerational gene silencing (49,52–56).

We describe a maternal effect in the transmission of the Eri phenotype and 22G sensor derepression and characterize the subjacent dynamics of sRNAs and mRNA targets (**Figs 1-3** and **S1-S3**). We show that both maternal and zygotic 26G-RNAs are sufficient for silencing. Absence of either the maternal or the zygotic pools can thus be compensated, enhancing the robustness of this system. We note, however, that sufficiency has only been tested with the described 22G sensor. It may be that the silencing of other targets has differential dependencies on maternal and zygotic 26G-RNA populations. The maternal effect rescue was observed for a variety of Eri genes, including *gtsf-1, rrf-3* and *ergo-1*, but not *alg-3/4*. Thus, these defects are related to impairment of sRNA populations directly associated with and downstream of ERGO-1. These results do not exclude a parental effect for ALG-3/4. In fact, a paternal effect on embryogenesis has been described for *rrf-3* mutants (30). Such phenotype most likely arises due to ALG-3/4 branch sRNAs.

Maternal rescue of Eri genes was previously reported (45), although the genetic basis for this phenomenon was not characterized further. We demonstrate that in the first Eri mutant generation, primary 26G-RNAs are downregulated, while their downstream 22G-RNAs are still present (**Fig 2**).

These ERGO-1-dependent 22G-RNAs, maintained in the absence of their primary triggers, seem to be competent to sustain silencing of ERGO-1 targets throughout life of the animal (**Fig 3**). Given that 1) ERGO-1 targets display higher expression during embryogenesis; and 2) upon disruption of endogenous RNAi by *rrf-3* mutation, targets become upregulated in all developmental stages (**Fig 3C**); maternally deposited ERGO-1-dependent factors may be especially required to initiate target silencing during embryogenesis and to prevent spurious expression throughout development. The ERGO-1-independent maintenance of this silencing response may be mechanistically similar to RNA-induced epigenetic silencing (RNAe), involving a self-perpetuating population of 22G-RNAs (49,57,58). Indeed, both processes depend on a nuclear Argonaute protein: HRDE-1 in RNAe (49,57,58) and NRDE-3 for ERGO-1-driven silencing (39,42,43). Self-perpetuating 22G-RNA signals may be also in place in the male germline (see below).

Our genetic experiments and sequencing data are fully consistent with maternal inheritance of 26G-RNAs. However, these may not be the only inherited agent. A non-mutually exclusive idea is that GTSF-1, as well as other ERIC proteins may be deposited in embryos to initiate production of zygotic sRNAs. In accordance with the latter, we have previously demonstrated that formation of the 26G-RNA generating ERIC is developmentally regulated (32). While in young adults there is a comparable amount of pre- and mature ERIC, in embryos there is proportionally more mature ERIC. These observations suggest that pre-ERIC might be deposited in the embryo to swiftly jumpstart zygotic 26G-RNA expression after fertilization.

### 26G-RNAs act as triggers to induce a self-sustained 22G-RNA-driven silencing in the male germline

We show that GTSF-1 is required in adult males to produce 26G- and downstream 22G-RNAs (**Fig 4**) analogous to its role in the hermaphrodite germline and in embryos (32). In addition, the bulk of targets from both 26G-RNA pathway branches seem to be deregulated. Interestingly, we note that although ERGO-1 and its cognate 26G-RNAs are not abundantly expressed in spermatogenic tissues (**Fig 4B**), *gtsf-1*-dependent, secondary 22G-RNAs mapping to these genes maintain gene silencing in the male germline (**Fig 4F-G**). In an analogous manner, we find that ALG-3/4 targets maintain 22G-RNAs in gravid adults (32), even though ALG-3/4 is not expressed at that stage. Mechanistically this may be closely related to how maternal 26G-RNAs can trigger 22G-RNA-driven silencing (see above). NRDE-3 is downstream of ERGO-1, likely silencing ERGO-1 targets throughout development. However, the Argonautes associated with 22G-RNAs mapping to 1) ERGO-1 targets in the male, and 2) to ALG-3/4 targets in gravid adults have not yet been identified.

### Two distinct ALG-3/4 regulatory mechanisms?

ALG-3/4-branch 26G-RNAs map very sharply to the 5’ and 3’ extremities of the targets, very close to the transcription start and end sites. We find that stronger targeting on the 3’ end does not drive robust gene silencing, and may even license expression, while targeting on the 5’ end is associated with stronger gene silencing. Targeting on the 3’ is consistent with RdRP recruitment to synthesize antisense secondary 22G-RNAs throughout the transcript. These may associate with CSR-1 and could have a positive effect on gene expression. The sharp 5’ peak in the metagene analysis could hint at additional regulatory modes, other than 22G-RNA targeting. 5’-end-bound ALG-3/4 could recruit other effector factors which promote RNA decay or translation inhibition, e.g. by inhibiting the assembly of ribosomes. Of note, when single targets are considered individually, 26G-RNA peaks at 5’ and 3’ can be simultaneously detected (**Figs 7, S2B** and **S5A**, left panels and **S6**). Hence, the resolution of a balance between Argonaute-sRNA complexes binding at 5’ and 3’ could determine regulatory outcome. Notably, we find shorter 3’ UTRs to be correlated with gene silencing (**Fig 6E**). In a model where predominant 3’ UTR targeting by Argonaute-sRNA complexes promotes gene expression, shorter 3’ UTRs and therefore less chance of sRNA binding may shift the balance towards gene silencing. Another possibility may be that longer 3’ UTRs contain binding sites for additional RNA binding proteins that may help to restrict RdRP activity on the transcript in question. Further work will be needed to test such ideas.

### An Argonaute negative feedback loop

In *C. elegans*, primary sRNAs trigger the production of secondary sRNAs in a feedforward loop. If left uncontrolled, such feedforward mechanisms can be detrimental to biological systems. Endogenous and exogenous RNAi pathways in *C. elegans* compete for limiting shared factors and the Eri phenotype is a result of such competition (33,44). Competition for shared factors is in itself a mechanism to limit accumulation of sRNAs. In support of this, exogenous RNAi was shown to affect endogenous sRNA populations, thus restricting the generations over which RNAi effects can be inherited (59).

We find that 26G-RNAs, likely ALG-3/4-bound, as well as 22G-RNAs map to *alg-3* and *alg-4* mRNAs (**Figs 7** and **S6**). In the absence of GTSF-1, a loss of these sRNAs is accompanied by a 2-3 fold upregulation of *alg-3* and *alg-4* on the mRNA level. This means that ALG-3 and ALG-4 may regulate their own expression. To the best of our knowledge, this is the first observation in *C. elegans* of Argonaute proteins regulating the expression of their own mRNA, and represents a very interesting case of a break on the positive feedback of the 26G-RNA pathway. Such regulation is not unprecedented. Complementary endo-siRNAs to *ago2* have been described in *Drosophila* S2 cells (60). Since AGO2 is required for the biogenesis and silencing function of endo-siRNAs, it is likely that Ago2 regulates itself in S2 cells.

Such direct self-regulation of Argonaute genes may constitute an important mechanism to limit RNAi-related responses, but the biological relevance of this regulation will need to be addressed experimentally. These observations do suggest that the Eri phenotype is but one manifestation of intricate cross-regulation governing the RNAi pathways of *C. elegans*.

## Materials and Methods

### *C. elegans* genetics and culture

*C. elegans* was cultured on OP50 bacteria according to standard laboratory conditions (61). Unless otherwise noted, worms were grown at 20°C. The Bristol strain N2 was used as the standard wild-type strain. All strains used and created in this study are listed in **S2 Table**.

### Microscopy

Wide-field photomicrographs were acquired using a Leica M165FC microscope with a Leica DFC450 C camera, and were processed using Leica LAS software and ImageJ.

### Genetic crosses using *dpy-4;gtsf-1* worms

#### Cross outline

We first linked *gtsf-1(xf43)* and *dpy-4(e1166)*. These genes are 2.62 cM apart, which does not comprise extremely tight linkage. Therefore, throughout the outcrossing scheme, worms were consistently genotyped for *gtsf-1* and phenotyped for *dpy-4*. We started by outcrossing *dpy-4;gtsf-1* hermaphrodites with N2 males (in a 1:2 ratio). *dpy-4(e1166)* is reported as being weakly semi-dominant (https://cgc.umn.edu/strain/CB1166). Indeed, heterozygote worms look only very slightly Dpy, therefore for simplicity, we refer to the heterozygote phenotype as “wild-type” throughout this work. Wild-type looking worms were selected in the F1 and F2 generations. The F2s were allowed to lay embryos for 1-2 days and then were genotyped for *gtsf-1(xf43)* using PCR. Progenies of non-recombined *gtsf-1* heterozygote worms were kept for follow up. F3 progenies that did not segregate *dpy* worms were discarded. F3 *dpys* were isolated, allowed to lay embryos, and genotyped for *gtsf-1(xf43)*. Progenies of non-homozygote mutant *gtsf-1(xf43)* worms were discarded.

#### RNAi

dsRNA against *lir-1* was supplemented to worms by feeding as described (62). L1 worms were transferred to RNAi plates and larval arrest was scored 2-3 days later. L1 F3 and F4 worms were transferred to RNAi plates blinded to genotype/phenotype (the *dpy* phenotype only shows clearly from L3 onwards).

#### Temperature-sensitive sterility assay

Single L1 F3 and F4 worms were transferred to OP50 plates, blinded to genotype/phenotype and grown at 25°C (the *dpy* phenotype only shows clearly from L3 onwards). Temperature sensitive-sterility was scored on the second day of adulthood and worms with unexpected genotype-phenotype were genotyped for *gtsf-1*.

#### RNA isolation

Approximately 550 hand-picked wild-type, Dpy F3 (referred in the text as mutant F1) and Dpy F4 (referred in the text as mutant F2) animals were used to isolate RNA (see cross description above, schematics in **Fig 2A**, and see below for RNA isolation protocol). Four independent outcrosses were performed and independent biological replicates (of wild-type, mutant F1 and mutant F2) were collected from each. Each sample was used to prepare small RNA and mRNA libraries (see below for details on library preparation).

### Growth and RNA isolation of adult males

*him-5(e1467)* and *him-5(e1467); gtsf-1(xf43)* worm populations were synchronized by bleaching, overnight hatching in M9 and plated on OP50 plates the next day. Worms were grown until adulthood for approximately 73 hours and 400-500 male animals were hand-picked for each sample, in biological triplicates, and used to isolate RNA (see below for RNA isolation protocol). Each sample was used to prepare small RNA and mRNA libraries (see below details on library preparation).

### Growth and RNA isolation of N2 and *rrf-3* worms

N2 and *rrf-3(pk1426)* animal populations were synchronized by bleaching, overnight hatching in M9 and plated on OP50 plates the next day. L1 animals were allowed to recover from starvation for 5 hours, and then were collected. L2 worms were collected 11 hours after plating. L3 animals were collected 28 hours after plating. L4 animals were collected 50 hours after plating, and young adults were collected 56 hours after plating. Embryo samples were collected from bleached gravid adult animals, followed by thorough washes with M9. Samples were collected in triplicate and RNA isolation proceeded as described below.

### RNA isolation

Worms were rinsed off plates and washed 4-6 times with M9 supplemented with 0.01% Tween. 50 µL of M9 plus worms were subsequently frozen in dry ice. For RNA isolation worm aliquots were thawed and 500 µL of Trizol LS (Life Technologies, 10296-028) was added and mixed vigorously. Next, we employed six freeze-thaw cycles to dissolve the worms: tubes were frozen in liquid nitrogen for 30 seconds, thawed in a 37°C water bath for 2 minutes, and mixed vigorously. Following the sixth freeze-thaw cycle, 1 volume of 100% ethanol was added to the samples and mixed vigorously. Then, we added these mixtures onto Direct-zol columns (Zymo Research, R2070) and manufacturer’s instructions were followed (in-column DNase I treatment was included).

### Library preparation for mRNA sequencing

NGS library prep was performed with Illumina‘s TruSeq stranded mRNA LT Sample Prep Kit following Illumina’s standard protocol (Part # 15031047 Rev. E). Starting amounts of RNA used for library preparation, as well as the number of PCR cycles used in amplification, are indicated in **S3 Table**. Libraries were profiled in a High Sensitivity DNA on a 2100 Bioanalyzer (Agilent technologies) and quantified using the Qubit dsDNA HS Assay Kit, in a Qubit 2.0 Fluorometer (Life technologies). Number of pooled samples, Flowcell, type of run and number of cycles used in the different experiments are all indicated in **S3 Table**.

### RppH treatment and library preparation for small RNA sequencing

For maternal effect sequencing, RNA was directly used for library preparation, or treated with RppH prior to library preparation. RppH treatment was performed as described in reference 47 with slight modifications. In short, 500 ng of RNA were incubated with 5 units of RppH and 10x NEB Buffer 2 for 1 hour at 37°C. Reaction was stopped by incubating the samples with 500 mM EDTA for 5 minutes at 65°C. RNA was reprecipitated in 100% Isopropanol and ressuspended in nuclease-free water. NGS library prep was performed with NEXTflex Small RNA-Seq Kit V3 following Step A to Step G of Bioo Scientific’s standard protocol (V16.06). Both directly cloned and RppH-treated libraries were prepared with a starting amount of 200ng and amplified in 16 PCR cycles. Amplified libraries were purified by running an 8% TBE gel and size-selected for 18 – 40 nts. Libraries were profiled in a High Sensitivity DNA on a 2100 Bioanalyzer (Agilent technologies) and quantified using the Qubit dsDNA HS Assay Kit, in a Qubit 2.0 Fluorometer (Life technologies). All 24 samples were pooled in equimolar ratio and sequenced on 1 NextSeq 500/550 High-Output Flowcell, SR for 1x 75 cycles plus 6 cycles for the index read.

RNA from adult males was RppH-treated as described above with the difference that 800 ng of RNA were used for RppH treatment. Library preparation of these samples was performed exactly as described above with the following modifications: starting amount of 460 ng; and amplification in 15 PCR cycles.

### Bioinformatic analysis

Sequencing statistics can be found in **S1 Table**

#### Small RNA read processing and mapping

Illumina adapters were removed with cutadapt v1.9 (63) (-a TGGAATTCTCGGGTGCCAAGG -O 5 -m 26 -M 38) and reads with low-quality calls were filtered out with fastq_quality_filter (-q 20 -p 100 -Q 33) from the FASTX-Toolkit v0.0.14. Using information from unique molecule identifiers (UMIs) added during library preparation, reads with the same sequence (including UMIs) were collapsed to remove putative PCR duplicates using a custom script. Prior to mapping, UMIs were trimmed (seqtk trimfq -b 4 -e 4) and reads shorter than 15 nucleotides (nts) were discarded (seqtk seq -L 15). Library quality was assessed with FastQC twice, for the raw and for the processed reads. Processed reads were aligned against the *C. elegans* genome assembly WBcel235 with bowtie v0.12.8 (64) (–tryhard –best –strata -v 0 -M 1). Reads mapping to structural genes were filtered out (r/t/s/sn/snoRNA) using Bedtools 2.25.0 (65) (bedtools intersect -v -s -f 0.9) and further analysis was performed using non-structural RNAs.

#### Small RNA class definition and quantification

Gene annotation was retrieved from Ensembl (release-38). Transposon coordinates were retrieved from wormbase (PRJNA13758.WS264) and added to the ensembl gene annotation to create a custom annotation used for further analysis. To define RNAs as belonging to particular classes of small RNA, mapped reads were categorized as follows: 21U-RNAs (piRNAs) are considered those sequences that are 21 nt long, and map sense to annotated piRNA loci; 22G-RNAs are those whose sequence is exactly 20-23 nts, have a guanine at their 5’ and map antisense to annotated protein-coding/pseudogenes/lincRNA/transposons; 26G-RNAs, are those which are 26 nt, and map antisense to annotated protein-coding/pseudogenes/lincRNA. Read filtering was done with a python script available at https://github.com/adomingues/filterReads/blob/master/filterReads/filterSmallRNAclasses.py which relies on pysam v0.8.1 an htslib wrapper (66), in combination with Bedtools intersect. Reads fulfilling these definitions were then counted for each library (total levels). Genome browser tracks were created using Bedtools (genomeCoverageBed -bg -split -scale -ibam -g), to summarize genome coverage normalized to mapped reads * 1 million (Reads Per Million or RPM), followed by bedGraphToBigWig to create the bigwig track. To quantify the effects on small RNAs of particular branches/pathways, we collected lists of genes previously identified as being targeted by these pathways: CSR-1 (36); NRDE-3 (43); Mutators (67); and WAGO-1 (52). ERGO-1 targets were defined as genes that lose oxidation-resistant 26G-RNAs (that are 3’ 2’-O-methylated) upon *gtsf-1* mutation (Table EV1, sheet 1.2 in reference 32). ALG-3/4 targets are defined as genes that lose 26G-RNAs upon *gtsf-1* mutation (Table EV1, sheet 1.1 in reference 32), excluding ERGO-1 targets. The genomic locations of 22G- and 26G-RNAs was then intersected with that of the genes, and counted for each library.

#### mRNA read processing and mapping

Library quality was assessed with FastQC before being aligned against the *C. elegans* genome assembly WBcel235 and a custom GTF, which included transposon coordinates (described above) with STAR v2.5.2b (–runMode alignReads – outSAMattributes Standard–outSJfilterReads Unique–outSAMunmapped Within–outReadsUnmapped None –outFilterMismatchNmax 2 –outFilterMultimapNmax 10 –alignIntronMin 21 –sjdbOverhang 79). Reads mapping to annotated features in the custom GTF were counted with subread featureCounts v1.5.1 (-s 2 -p -F GTF –donotsort -t exon -g gene_id). Coverage tracks were generated with deepTools v2.4.3 (bamCoverage –smoothLength 60 –binSize 20 – normalizeUsingRPKM) (68).

#### Differential expression/small RNA targeting

Reads mapping to annotated features in the custom GTF were counted with htseq-count v0.9.0 (69)(htseq-count -f bam -m intersection-nonempty -s reverse) for sRNA-seq data, and with subread featureCounts v1.5.1 (70) (-s 2 -p -F GTF – donotsort -t exon -g gene_id) for mRNA-seq. Pairwise differential expression comparisons were performed with DESeq2 v.1.18.1 (71). For the selection of genes differentially targeted (sRNA) or expressed (mRNA), a further cut-off of at least a 1.5 fold-change difference between conditions was applied. As previously reported (32), due to the observed global depletion of 26G-RNA reads in some samples (sRNA), DESeq2 library sizes computed from all reads 18-30 nt in each sample were for 26G-RNA differential analyses. Gene expression in RPKM (Reads Per Kilobase Million) was calculated by retrieving the fragments/counts per million mapped fragments from the DEseq2 object (fpm(object, robust = TRUE)) and normalizing to gene length.

#### Metagene analysis

The average coverage at each gene from a particular branch was determined with deepTools v2.4.3 (computeMatrix scale-regions –metagene –missingDataAsZero –b 250 -a 250 –regionBodyLength 2000 –binSize 50 –averageTypeBins median), using the transcript locations of each gene, and plotted with plotProfile –plotType se –averageType mean –perGroup.

#### UTR targeting by 26G-RNAs

To identify genes predominantly targeted at their 5’ or 3’, coverage values of scaled genes were obtained with deepTools, as done for the metagene analysis (see above), with the difference that only the WT track was used, and options -a and -b were set to 0. That is, only the scaled body regions were used. 5’ and 3’ sRNA targeting was defined for each gene based on the coverage at the first or last 25% of the scaled gene body. The genes were then classified in low, medium or high targeting if they were in the 0-25, 25-75, or 75-100 percentile of the sRNA coverage distribution for either the 5’ or the 3’. Primarily 5’ or 3’ targeted genes were further defined if they were in the 5’ high and 3’ low category (5’ targeted), or high in the 3’ and low in the 5’ (3’ targeted).

### Accession numbers

All sequencing data has been submitted to SRA, accession number PRJNA497368.

## Author Contributions

Conceptualization, M.V.A. and R.F.K.; Investigation, M.V.A.; Formal Analysis, M.V.A., A.M.dJ.D.; Visualization, M.V.A., A.M.dJ.D.; Writing – Original Draft, all authors contributed; Writing – Review & Editing, all authors contributed; Funding acquisition, R.F.K.

## Acknowledgements

We thank all the members of the Ketting lab for great help and discussion. A special thanks to Yasmin el Sherif and Svenja Hellmann for excellent technical assistance and to Jan Schreier for producing the *mut-16(xf142)* mutant. The authors are grateful to Hanna Lukas, Clara Werner and Maria Mendez-Lago of the IMB genomics core facility for library preparation. We thank the IMB Media Lab for consumables. We also acknowledge the *Caenorhabditis* Genetics Center (CGC), which is funded by NIH Office of Research Infrastructure Programs (P40 OD010440), for providing worm strains. This work was supported by a Deutsche Forschungsgemeinschaft grant KE 1888/1-1 (Project Funding Programme to R.F.K.).

## Supporting Information Captions

**Fig S1.**
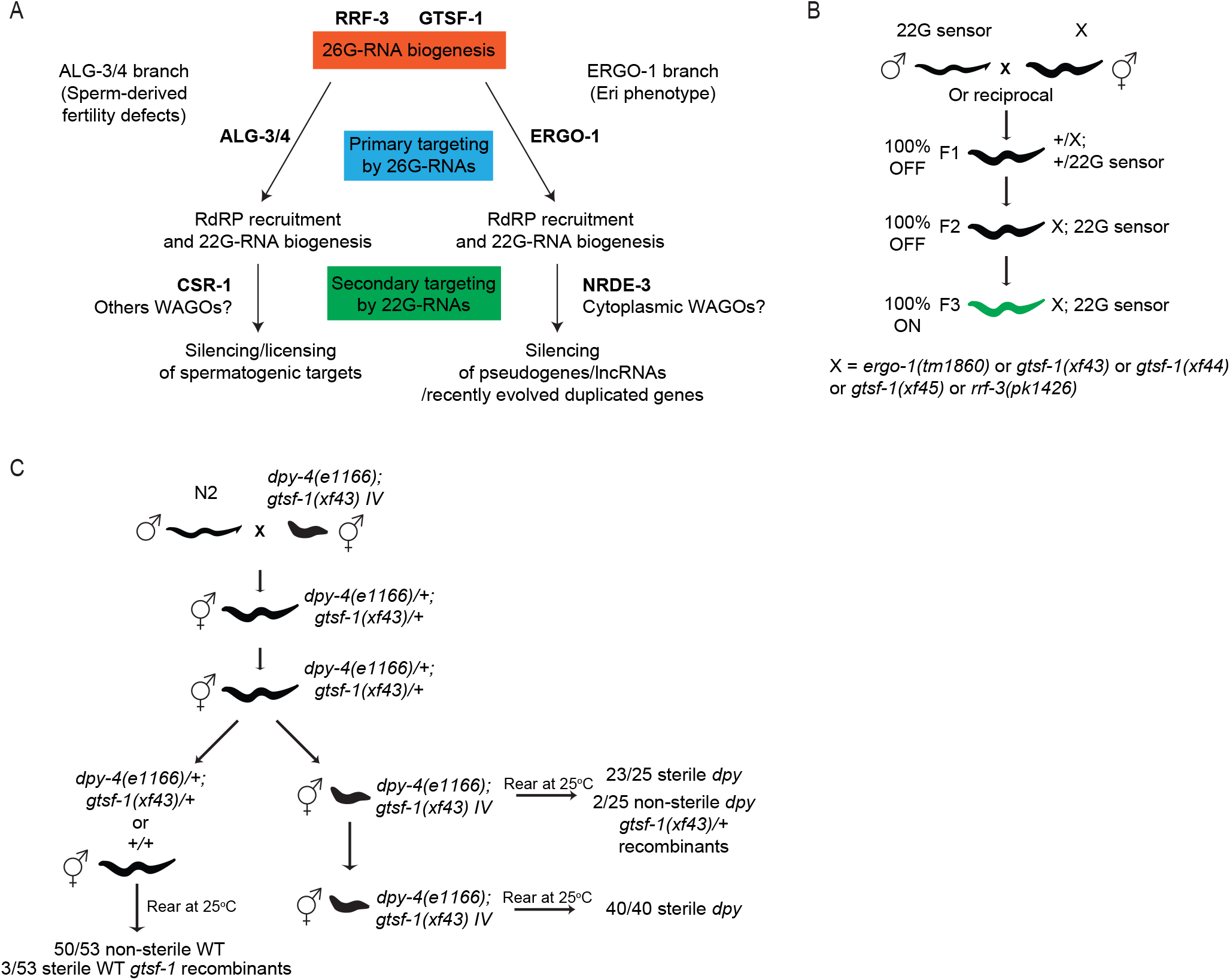
**Parental effects of the 26G-RNA pathway**. (A) Illustration of the current understanding of 26G-RNA pathways. 26G-RNAs are produced by RRF-3, assisted by GTSF-1 and other accessory factors. 26G-RNAs can associate with ALG-3/4 in the spermatogenic gonad (ALG-3/4 branch) or with ERGO-1 in oocytes and embryos (ERGO-1 branch). Upon target binding, RdRPs are recruited and synthesize secondary 22G-RNAs. NRDE-3 binds ERGO-1 branch 22G-RNAs, while is downstream of ALG-3/4 branch 26G-RNAs. Other unidentified Argonautes may play a role in these pathways. (B) Schematics of genetic crosses of mutant strains with the 22G sensor. Green worms illustrate derepression of the 22G sensor. “X” corresponds to different mutant alleles that share the same maternal rescue. (C) Experimental setup to address the maternal transmission of the temperature-sensitive sterility phenotype at 25°C. Worms were constantly grown at 20°C until transfer to 25°C to assay sterility. L2-L3 worms were transferred to 25°C.

**Fig S2.**
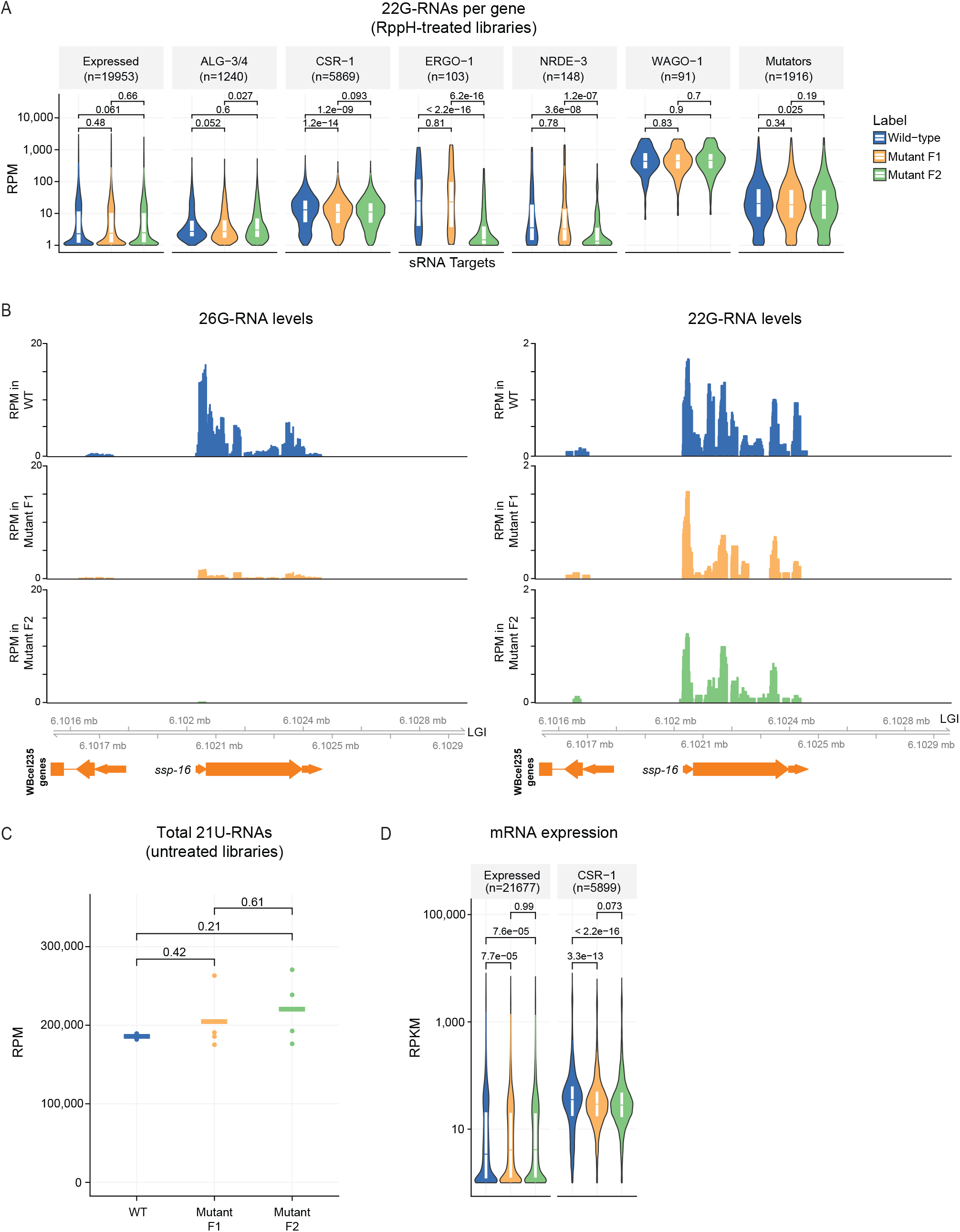
**Dynamics of RNA expression upon *gtsf-1* mutation**. (A) Violin plot showing the distribution of 22G-RNAs mapping, per gene, to known targets of diverse sRNA pathways. (B) Genome browser tracks of *ssp-16*, a known ALG-3/4 target, showing mapped 26G- and 22G-RNAs. 26G- and 22G-RNA tracks were obtained from untreated and RppH-treated libraries, respectively. (C) Total 21U-RNA levels in different generations/phenotype, in RPM. (D) Distribution of normalized mRNA expression of CSR-1 targets in RPKM. Violin plots in (A) and (D) show the distribution density of the underlying data. The top and bottom of the embedded box represent the 75th and the 25th percentile of the distribution, respectively. The line in the box represents the median. P-values were calculated with a two-sided unpaired Mann-Whitney/Wilcoxon rank-sum test.

**Fig S3.**
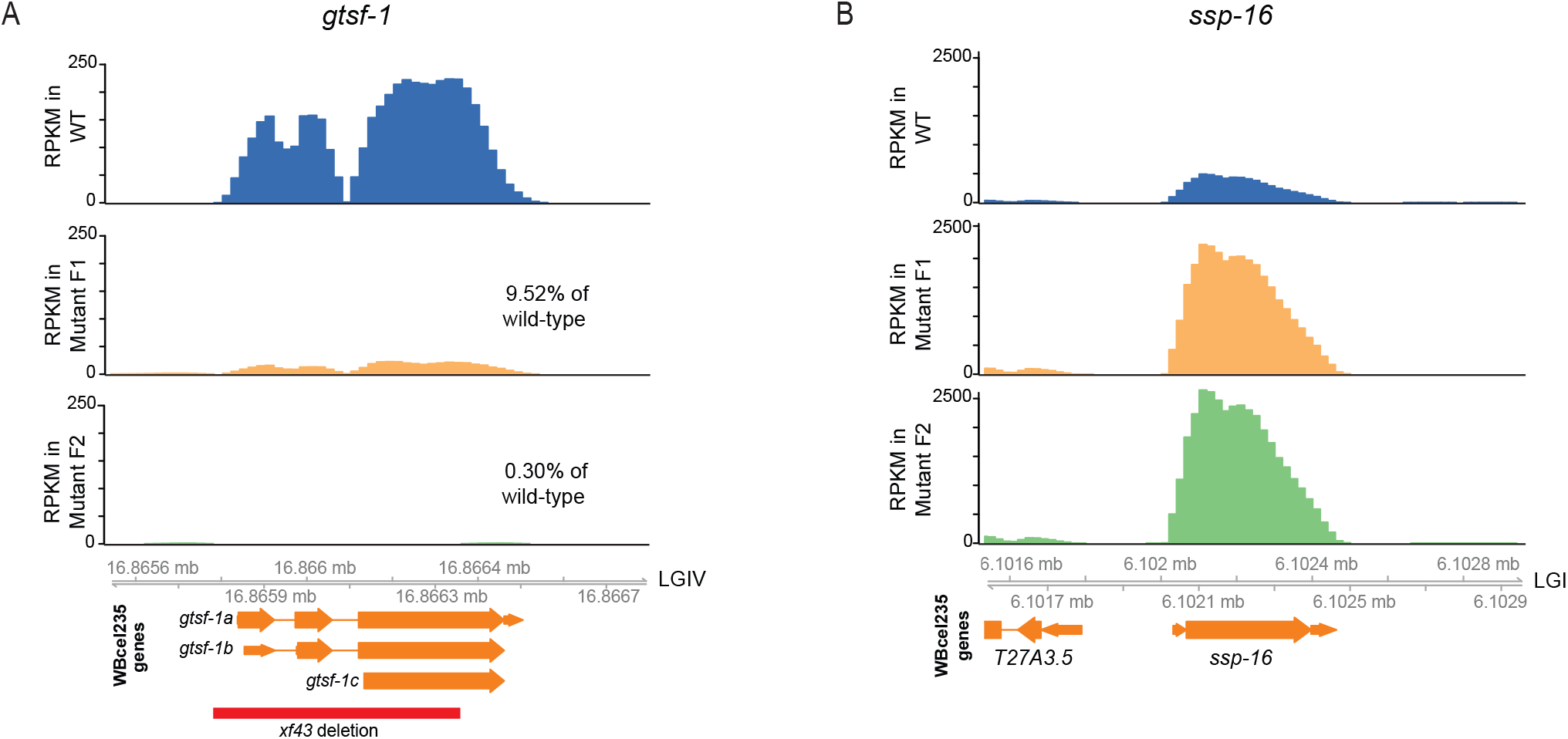
**mRNA dynamics upon *gtsf-1* mutation**. (A) Genome browser tracks displaying *gtsf-1* mRNA levels in RPKM. The *gtsf-1(xf43)* deletion allele is represented below. *gtsf-1* levels in the mutant F1 cover the *xf43* deletion sequence, thereby indicating contamination with Dpy worms that recombined a wild-type copy of *gtsf-1*. The mutant F2 was isolated from mutant F1 Dpy whose *gtsf-1* genotype was confirmed. Therefore, as expected, the only observed reads are flanking the *xf43* deletion. (B) Genome browser tracks with the mRNA levels, in RPKM, of *ssp-16*. Upregulation occurs immediately in the F1, indicating no maternal effect.

**Fig S4.**
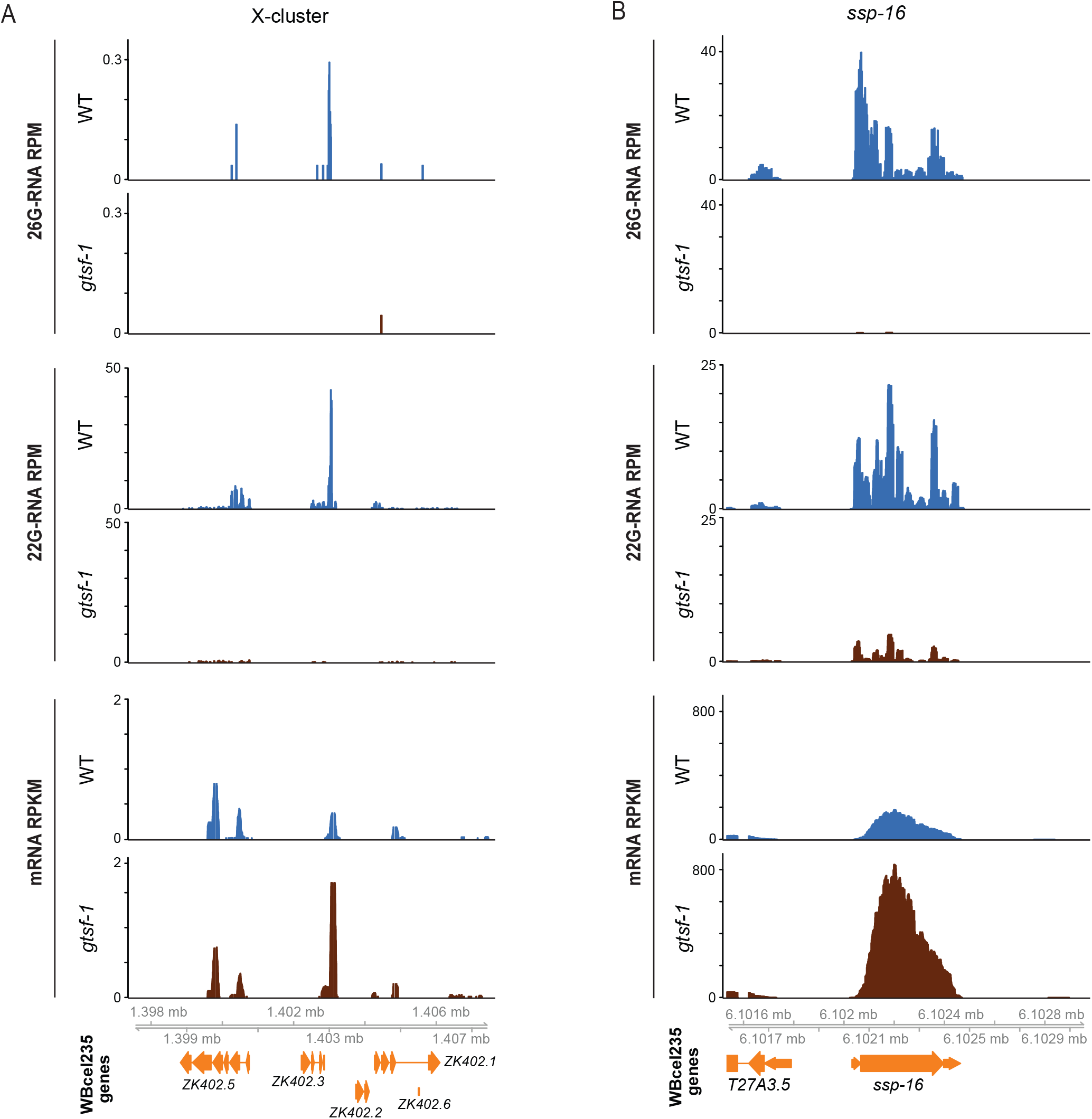
**sRNA and mRNA profiles of ERGO-1 and ALG-3/4 targets in males**. (A-B) RPM levels of 26G-RNAs (upper panels) and 22G-RNAs (middle panels) mapping to the X-cluster (A) and *ssp-16* (B). Lower panels show RPKM mRNA levels of these targets. WT, wild-type.

**Fig S5.**
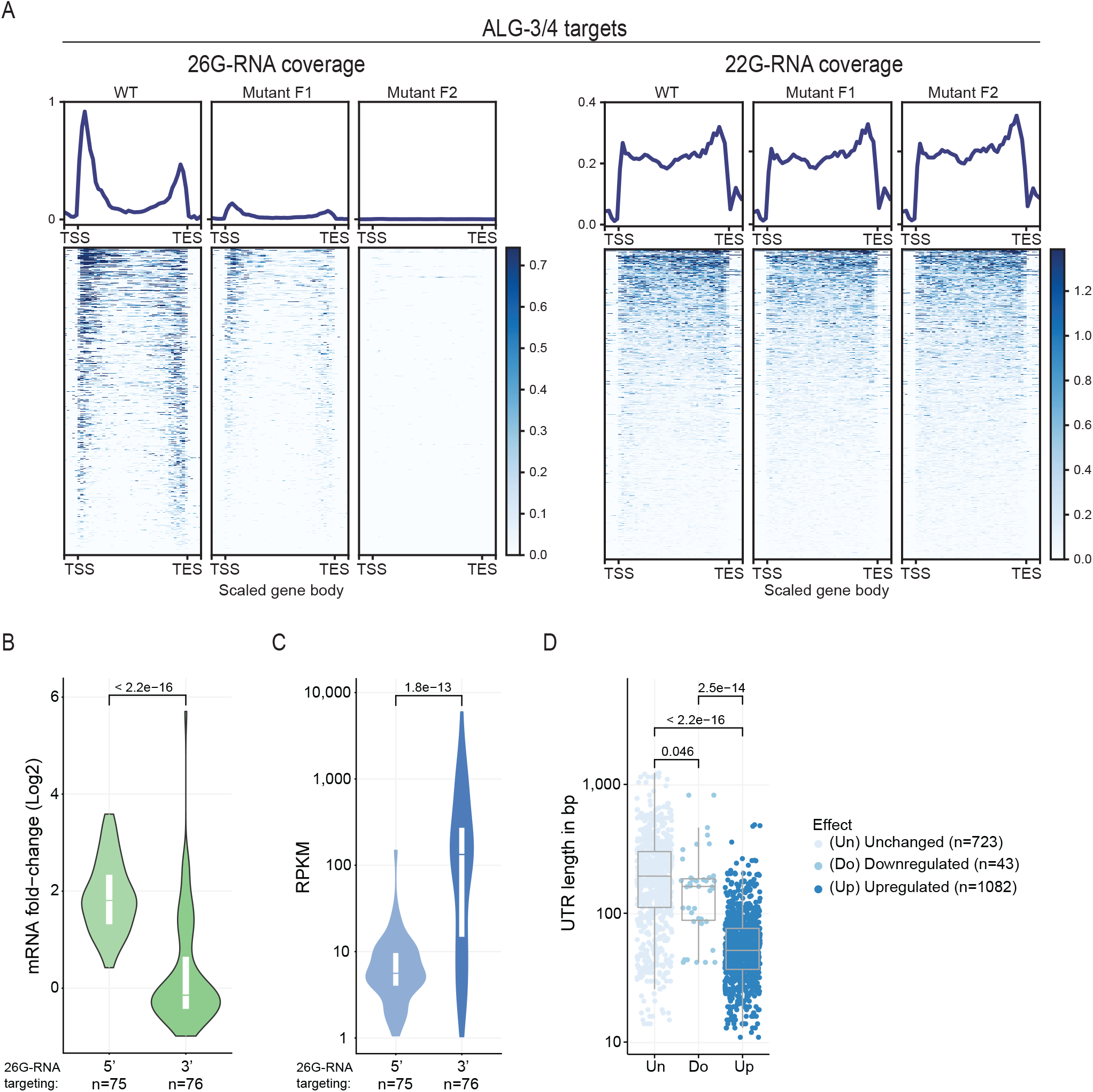
**Predictors of regulatory outcome by ALG-3/4 in young adults**. (A) Metagene analysis of 26G-(left panel) and 22G-RNAs (right panel) mapping to ALG-3/4 targets in young adult datasets, from our maternal effect setup (as in **Fig 2A**). On the upper part of each panel is the mean coverage profile for sRNA species in every generation. On the lower part of each panel, the heatmaps show the density across individual targets. Target gene body length was scaled between transcription start site (TSS) and transcription end site (TES). Moreover, the regions comprising 250 nucleotides immediately upstream of the TSS and downstream of the TES are also included. Simultaneous 26G-RNA targeting in the 5’ and 3’ can be observed in some genes. (B) Violin plot depicting the regulation of ALG-3/4 target genes predominantly targeted on the 5’ or on the 3’ by 26G-RNAs. (C) Violin plot showing the wild-type expression levels of ALG-3/4 target genes predominantly targeted on the 5’ on or the 3’ by 26G-RNAs. (D) 3’ UTR lengths of all the transcript isoforms annotated for ALG-3/4 target genes, according to effect on gene expression. All the panels of this Fig were prepared using young adult sequencing datasets from our maternal effect experiments. In B and D, regulatory outcome was defined as differential gene expression between the wild-type and *gtsf-1* mutant F2. For (B-D) we used male sequencing datasets. Violin plots in (B-C) and the boxplot in (D) show the distribution of the data. The top and bottom of the embedded boxes represent the 75th and the 25th percentile of the distribution, respectively. The lines in the boxes represent the median. P-values were calculated with a two-sided unpaired Mann-Whitney/Wilcoxon rank-sum test.

**Fig S6.**
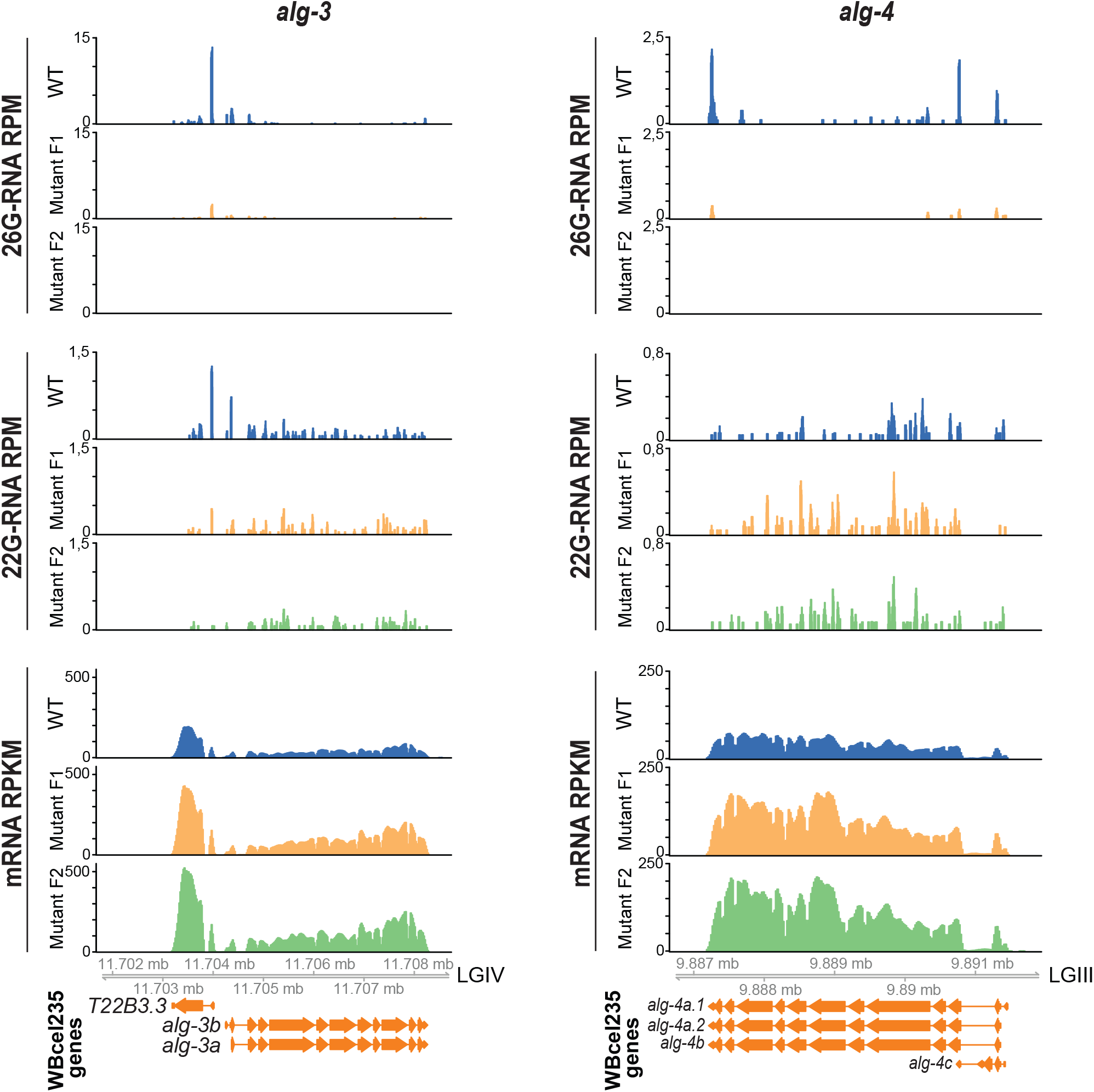
**ALG-3 and ALG-4 are engaged in a negative feedback loop in young adults**. Genome browser tracks showing 26G-RNAs (upper panels) and 22G-RNAs (middle panels) mapping to *alg-3* (left panels) and *alg-4* (right panels), in RPM. Lower panels show the RPKM mRNA levels of *alg-3* (on the left) and *alg-4* (on the right). Sequencing datasets of young adults from our maternal effect setup were used.

**S1 Table. Sequencing statistics**.

**S2 Table. Strains used in this study**.

**S3 Table. Specifics of library preparation and sequencing**.

